# VARP binds SNX27 to promote endosomal supercomplex formation on membranes

**DOI:** 10.1101/2024.07.11.603126

**Authors:** Mintu Chandra, Amy K. Kendall, Marijn G. J. Ford, Lauren P. Jackson

## Abstract

Multiple essential membrane trafficking pathways converge at endosomes to maintain cellular homeostasis by sorting critical transmembrane cargo proteins to the plasma membrane or the *trans*-Golgi network (TGN). The Retromer heterotrimer (VPS26/VPS35/VPS29 subunits) binds multiple sorting nexin (SNX) proteins on endosomal membranes, but molecular mechanisms regarding formation and regulation of metazoan SNX/Retromer complexes have been elusive. Here, we combine biochemical and biophysical approaches with AlphaFold2 Multimer modeling to identify a direct interaction between the VARP N-terminus and SNX27 PDZ domain. VARP and SNX27 interact with high nanomolar affinity using the binding pocket established for PDZ binding motif (PDZbm) cargo. Specific point mutations in VARP abrogate the interaction *in vitro*. We further establish a full biochemical reconstitution system using purified mammalian proteins to directly and systematically test whether multiple endosomal coat complexes are recruited to membranes to generate tubules. We successfully use purified coat components to demonstrate which combinations of Retromer with SNX27, ESCPE-1 (SNX2/SNX6), or both complexes can remodel membranes containing physiological cargo motifs and phospholipid composition. SNX27, alone and with Retromer, induces tubule formation in the presence of PI(3)*P* and PDZ cargo motifs. ESCPE-1 deforms membranes enriched with Folch I and CI-MPR cargo motifs, but surprisingly does not recruit Retromer. Finally, we find VARP is required to reconstitute a proposed endosomal “supercomplex” containing SNX27, ESCPE-1, and Retromer on PI(3)*P*-enriched membranes. These data suggest VARP functions as a key regulator in metazoans to promote cargo sorting out of endosomes.

## Introduction

Large multi-subunit coat protein complexes initiate distinct trafficking pathways by forming hubs at organellar membranes (*1*). Coats recognize and cluster specific lipid and transmembrane protein cargo, such as receptors, channels, or enzymes, for packaging into vesicles or tubules. Coats also recruit machinery required to form vesicles or tubules that will ultimately break off from the donor membrane to deliver cargo. Specific coats have traditionally been thought to define different trafficking pathways to ensure transmembrane cargoes are directed to the correct destination in a timely manner (*2–5*). On endosomal membranes, the Retromer heterotrimer composed of VPS26, VPS29, and VPS35 subunits (*6*, *7*) (Figure S1A) plays a role in sorting many cargoes. Retromer can directly bind multiple SNXs to form endosomal coat complexes (*4*, *8–16*). In yeast, Retromer exists as a stable pentamer composed of the Vps26/Vps35/Vps29 heterotrimer with a sorting nexin (SNX) heterodimer, Vps5 and Vps17 (*4*, *6*, *14*, *15*, *17*). In metazoans, Retromer has apparently expanded its repertoire to interact with additional SNXs, including orthologous SNX-BAR heterodimers (SNX1/SNX5, SNX2/SNX6); SNX3; and metazoan-specific SNX27 (*4*, *10*, *12*, *15*, *16*, *18–23*). Disruption of Retromer and SNX-mediated trafficking pathways through mutations or protein loss are associated with carboxypeptidase (CPY) mis-sorting in yeast (*7*) and various human neurologic disorders including Alzheimer’s disease, Parkinson’s disease, and Down’s syndrome (*24*, *25*).

Sorting nexin proteins belong to a large protein family defined by the presence of a Phox Homology (PX) domain, which recognizes membranes enriched with phosphoinositides (*26–28*). Specific SNX proteins have been shown to act as cargo adaptors by trafficking key proteins from endosomes to the plasma membrane or to the *trans*-Golgi complex (*8*, *28*, *29*). A subset of SNX family members have been shown to deform membranes (*20*, *30–33*). BAR domains (Bin/Amphiphysin/Rvs) found in SNX1 and SNX2 have previously been shown to form homodimers (*30*, *33*) to drive or stabilize membrane curvature through a scaffolding mechanism (*18*, *34*, *35*). In some cases, BAR domains can deform membranes through a second mechanism by using an amphipathic helix to insert into one leaflet to generate asymmetry and drive membrane curvature (*30*, *32*, *36*, *37*).

In mammals, Retromer is thought to associate with specific SNX-BAR heterodimers (SNX1/SNX5 or SNX2/SNX6) to retrieve cargo from endosomes back to the TGN (*18*, *22*, *31*, *32*, *34*) in a pathway analogous to the yeast pentamer. In yeast, the Vps5/Vps17 SNX-BAR heterodimer is proposed to be the functional homolog of the mammalian SNX1/5 or SNX2/6 complexes (*8*, *35*). SNX1/2 and SNX5/6 arose from gene duplication events (*11*, *32*, *38*, *39*), and the purpose of retaining two heterodimer complexes in metazoan cells remains unclear. More recently, SNX1/SNX5 or SNX2/SNX6 heterodimers have been proposed to form the Endosomal SNX–BAR Sorting Complex for Promoting Exit 1 (ESCPE-1) complex (*22*, *33–35*). In this model, SNX1 or SNX2 are proposed to act in membrane deformation while SNX5 or SNX6 further contribute to curvature and recognize specific motifs found in cargo, including CI-MPR (*22*, *34*, *40*) (Figure S1B). Another protein, SNX3, is conserved from yeast to humans and implicated in sorting distinct cargoes including DMT1-II and Wntless (*41*). SNX3 lacks a BAR domain, or any other module known to impact membrane bending. In recent years, pioneering structural studies revealed how some SNX proteins assemble with Retromer on membranes using cryo-ET (*19*, *20*, *41–45*). These studies demonstrated tubular structures with Retromer forming V-shaped arches on top of various SNX proteins (yeast Vps5 homodimer; yeast SNX3; or mammalian SNX3) (*20*, *42*, *46*), although these structures use truncated SNX proteins lacking N-termini for technical reasons.

The final SNX protein implicated in Retromer-mediated sorting is SNX27, which is unique to metazoans and required for recycling hundreds of transmembrane protein receptors. SNX27 (Figure S1C) possesses a different domain architecture as compared to SNX-BARs or SNX3. The SNX27 N-terminal PDZ (postsynaptic density 95/discs large/zonula occludens-1) domain binds transmembrane proteins with PDZ binding motifs (PDZbms) having the sequence T/S-X-Φ, and PDZbm cargo binding is enhanced by the Retromer VPS26 subunit. The central PX domain enables membrane recruitment through its affinity for PI(3)*P*. The C-terminal FERM (band 4.1/ezrin/radixin/moesin) domain is an interaction module proposed to undertake multiple functions. The FERM directly binds short motifs found in SNX1 and SNX2 N-termini (*22*, *40*, *47*), as well as small Ras GTPases (*48*, *49*) and transmembrane proteins with NPxY motifs (*12*, *26*, *27*, *40*, *48–53*). In metazoans, SNX27/Retromer is proposed to recycle specific cargoes from endosomes to the plasma membrane (*8*, *15*, *22*, *50*, *54*, *55*). SNX27/Retromer has been biochemically and functionally linked to SNX-BARs through binding the N-terminus of SNX1 and SNX2 (*22*, *40*, *47*). Overall, current models in metazoans suggest different combinations of SNX proteins bind Retromer to promote either retrieval or recycling of specific cargoes. However, there are currently no published data to demonstrate whether SNX27/Retromer can deform membranes as yeast SNX-BAR/Retromer or SNX3/Retromer complexes can.

In addition to SNX proteins, metazoan Retromer interacts with multiple accessory proteins that regulate its role in endosomal trafficking. Important examples include VARP (also known as ANKRD27; Figure S1D), Rab7, TBC1D5, and the WASH complex subunit FAM21 (*8*, *56–62*). Among these proteins, VARP has emerged as a key player in regulating late endosomal dynamics (*54*, *61–65*). VARP (VPS9 domain Ankyrin Repeat Protein) is a multi-domain protein with a VPS9 domain and two ankyrin repeat domains serving as protein-protein interaction modules. VARP functions as a Rab32/38 effector, and it displays GEF activity towards Rab21. VARP directly binds VAMP7, an R-SNARE involved in endocytic and secretory pathways (*54*, *61–65*). VARP recruitment to endosomal membranes relies on binding to VPS29 using two conserved cysteine-rich motifs located adjacent to the two ankyrin repeat domains (*54*, *61–65*). A mass spectrometry (MS)-based proteomics study of the SNX27 interactome (*66*) recently revealed VARP as a potential SNX27 interacting partner, but the biochemical basis for direct binding between SNX27 and VARP has not been established.

In this study, we use biochemical and biophysical methods together with AlphaFold Multimer modeling to demonstrate a new molecular interaction between the N-terminal folded domain of VARP and the SNX27 PDZ domain via the well-established PDZbm binding pocket. Biochemical pulldown assays confirm both full-length proteins and individual domains interact, while biolayer interferometry (BLI) establishes a relatively strong trafficking interaction (high nM K_D_). Structure-based point mutations generated based on AlphaFold computational structures further define sequence requirements for the interaction. Next, we developed a biochemical reconstitution system using purified proteins to systematically establish which combinations of endosomal coat proteins are recruited to liposomes in the presence of relevant phospholipids and cargo motifs. We paired liposome pelleting assays with negative stain electron microscopy (EM) to ascertain conditions under which combinations of SNX and Retromer proteins can bind and tubulate membranes. These experiments demonstrate for the first time how metazoan SNX27 on its own and together with Retromer can deform and tubulate membranes enriched with PI(3)*P* and PDZ cargo motifs. We further show how an ESCPE-1 complex containing the SNX2/SNX6 heterodimer can deform and tubulate membranes enriched with Folch I and CI-MPR cargo motifs, but it does not recruit Retromer. These two different endosomal coats yield tubules having different physical diameters. Finally, we tested whether ESCPE-1 can engage SNX27/Retromer to form the proposed endosomal “supercomplex” (*18*, *21*, *22*) and find supercomplex formation depends on the presence of VARP in the reconstitution system. The VARP N-terminus alone is sufficient to promote supercomplex formation *in vitro*, and structure-guided VARP point mutations abrogate the interaction on liposome membranes. Together, these results reveal an important new role for VARP in regulating endosomal trafficking and advance our understanding of how different endosomal coat complexes generate distinct carriers for efficient cargo sorting out of endosomes.

## Results

### VARP directly binds SNX27 *in vitro*

Over the past decade, numerous VARP protein binding partners have been identified (Figure S1D), highlighting its diverse roles in Retromer-mediated endosomal trafficking pathways. The SNX27 interactome has been explored using proteomics approaches (*66*), which suggest VARP and SNX27 may directly bind each other. We tested whether SNX27 could directly bind VARP using recombinant purified proteins in pulldown experiments (Figure 1A). Glutathione-S-transferase (GST)-tagged full-length SNX27 (GST-SNX27) was used as bait and full-length VARP with a C-terminal 10xHis tag (VARP-H10) as prey. For these experiments, we expressed and purified full-length human VARP in a mammalian expression system (see Methods). GST-SNX27 efficiently pulls down VARP-H10 (Figure 1A); the interaction is detected on a Coomassie stained gel and verified using an antibody against the 10xHis tag on VARP. The interaction between full-length VARP and SNX27 proteins was further quantified using biolayer interferometry (BLI). A robust dose-dependent increase in the binding between SNX27 and VARP was observed; the calculated average binding affinity (K_D_) is sub-micromolar (0.34 ± 0.01 µM) (Figure 1B; Table 1) with 1:1 stoichiometry (see Methods). As a positive control, we also measured full-length VARP with Retromer (Figure 1C) at a range of concentrations. These data reveal a nanomolar binding affinity (K_D_ = 0.07 ± 0.01 µM) (Figure 1C; Table 1) and 1:2 stoichiometry between one VARP and two Retromer complexes (see Methods), in line with published data (*63*).

**Figure 1.**
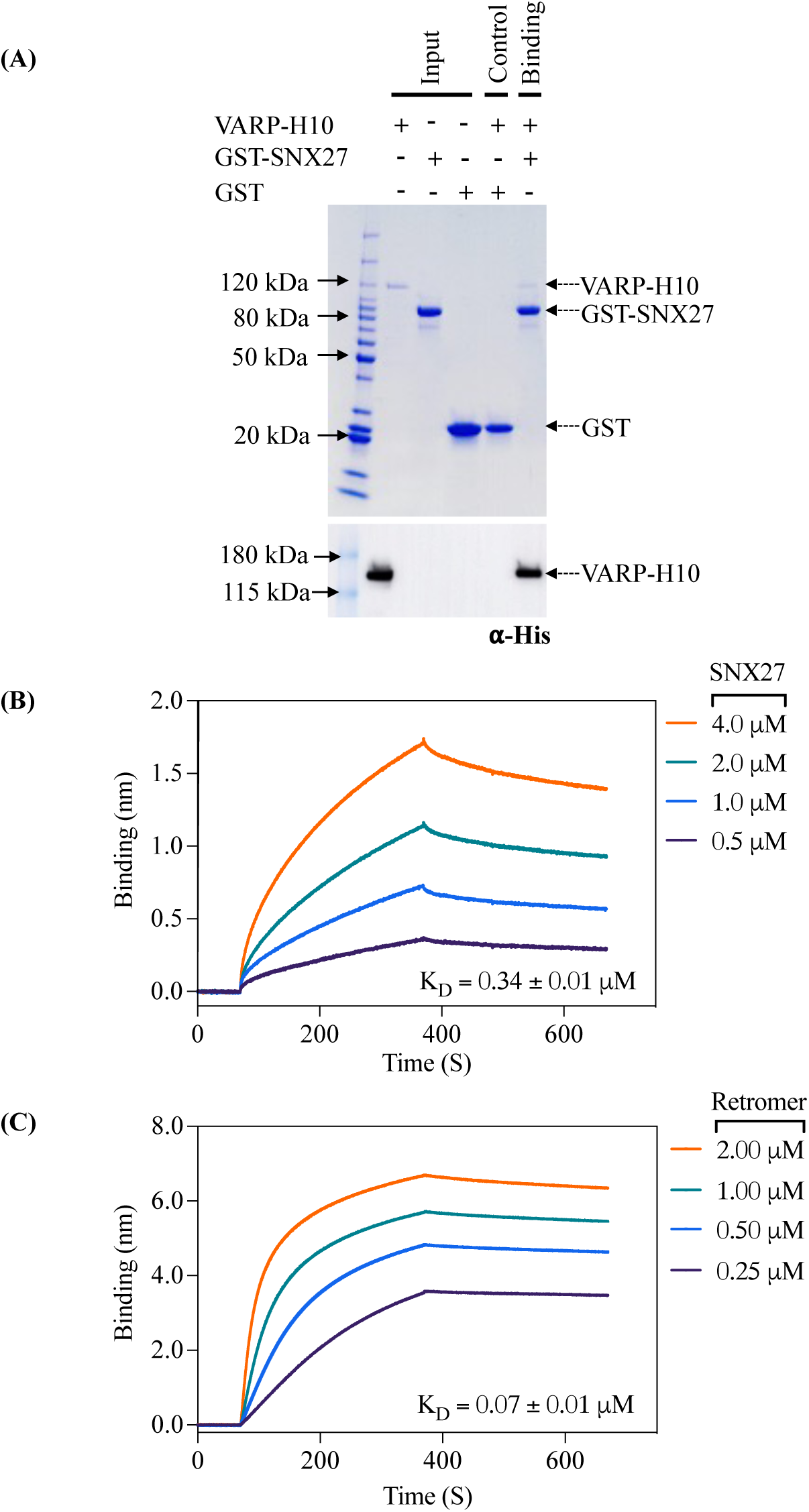
VARP directly binds SNX27 *in vitro*. **(A)** Coomassie Blue stained SDS-PAGE gel showing GST pulldown experiments with purified GST-SNX27 as bait and VARP-H10 as prey (top panel) and ⍺-His blot (bottom panel). GST-SNX27 is sufficient to pull down full-length VARP *in vitro*. **(B and C)** Biophysical data using the BLI Octet system demonstrate VARP-H10 directly binds SNX27 **(B)** or Retromer **(C)** with low micromolar affinity with K_D_ values below 1 μM. VARP-H10 was loaded onto a Ni-NTA biosensor, and data were obtained for either SNX27 or Retromer (positive control) at a range of concentrations. Fitted data were plotted in GraphPad Prism, and binding kinetics were calculated using the Octet R8 analysis software package.

**Table 1.** Kinetics and thermodynamic BLI measurements of VARP with SNX27 and Retromer. Data were analyzed using the Octet R8 analysis software package (Sartorius).

| **Loading Protein** | **Loading Protein Conc. (mg/ml)** | **Sample Protein** | **Sample Protein Conc. (μM)** | **K<sub>D</sub> (μM)** | **k<sub>a1</sub> (1/μMs)** | **k<sub>a2</sub> (1/μMs)** | **k<sub>dis1</sub> (1/s)** | **k<sub>dis2</sub> (1/s)** | **RSS** | **R<sup>2</sup>** | **Binding Model** |
| --- | --- | --- | --- | --- | --- | --- | --- | --- | --- | --- | --- |
| VARP FL | 0.05 | Retromer | 0.25 – 2 | 0.07 ± 0.01 | 5.04*10<sup>-3</sup> ± 1.88*10<sup>-5</sup> | 5.35*10<sup>-8</sup> ± 7.63*10<sup>-9</sup> | 3.63*10<sup>-4</sup> ± 5.77*10<sup>-6</sup> | 8.91*10<sup>-3</sup> ± 9.33*10<sup>-4</sup> | 3.33 | 0.99 | 1:2 |
| VARP FL | 0.05 | SNX27 FL | 0.5 – 4 | 0.34 ± 0.03 | 2.24*10<sup>-3</sup> ± 9.91*10<sup>-6</sup> | N.A. | 7.54*10<sup>-4</sup> ± 4.73*10<sup>-6</sup> | N.A. | 16.87 | 0.99 | 1:1 |
| N-VARP | 0.05 | SNX27 FL | 0.25 – 2 | 0.17 ± 0.02 | 3.21*10<sup>-3</sup> ± 2.38*10<sup>1</sup> | N.A. | 5.68*10<sup>-4</sup> ± 7.99*10<sup>-6</sup> | N.A. | 2.75 | 0.95 | 1:1 |
| N-VARP | 0.05 | SNX27 PDZ | 0.25 – 2 | 0.14 ± 0.03 | 4.31*10<sup>-3</sup> ± 4.15*10<sup>1</sup> | N.A. | 6.42*10<sup>-4</sup> ± 1.35*10<sup>-5</sup> | N.A. | 0.95 | 0.87 | 1:1 |
| N-VARP | 0.05 | SNX27 PX | 0.25 – 2 | N.B. | N.B. | N.A. | N.B. | N.A. | N.B. | N.B. | N.B. |
| N-VARP | 0.05 | SNX27 FERM | 0.25 – 2 | N.B. | N.B. | N.A. | N.B. | N.A. | N.B. | N.B. | N.B. |

### The folded VARP N-terminal domain directly binds the SNX27 PDZ domain

We next turned to AlphaFold Multimer version 2.3 (AF2.3) to generate computational models for the interaction between full-length SNX27 (SNX27 FL) and full-length VARP (VARP FL) (Figure S2). In line with biochemical (Figures 1A) and biophysical (Figure 1B) data, AF2.3 models indicated the N-terminal globular domain of VARP (N-VARP) specifically engages the SNX27 PDZ domain (Figure 2A). We next generated models using only the VARP N-terminus and SNX27 PDZ domains. These runs consistently produced highly reproducible models with high-confidence pLDDT scores (approaching 90; Figure S3) and ipTM scores (close to 0.9), along with low PAE scores (close to 0). These metrics strongly support a binding interface between the SNX27 PDZ domain and N-VARP (Figure 2A). The PISA server (*67*) was used to independently analyze the predicted interface between the SNX27 PDZ and N-VARP. PISA analysis reports a substantial buried surface area (1506.4 Å²), which further supports a biological interaction leading to formation of a stable complex (Figure S4A, S4C). We further evaluated AF2.3 model geometry using MolProbity (*68*) (Table 2). MolProbity reports favorable rotamers and Ramachandran values as well as low Clashscore. Data quality for the AF2.3 model reported here are in line with experimental data from an experimental X-ray structure of the SNX27 PDZ domain with a PDZbm cargo motif (PDB ID: 5EM9; Table 2).

**Figure 2.**
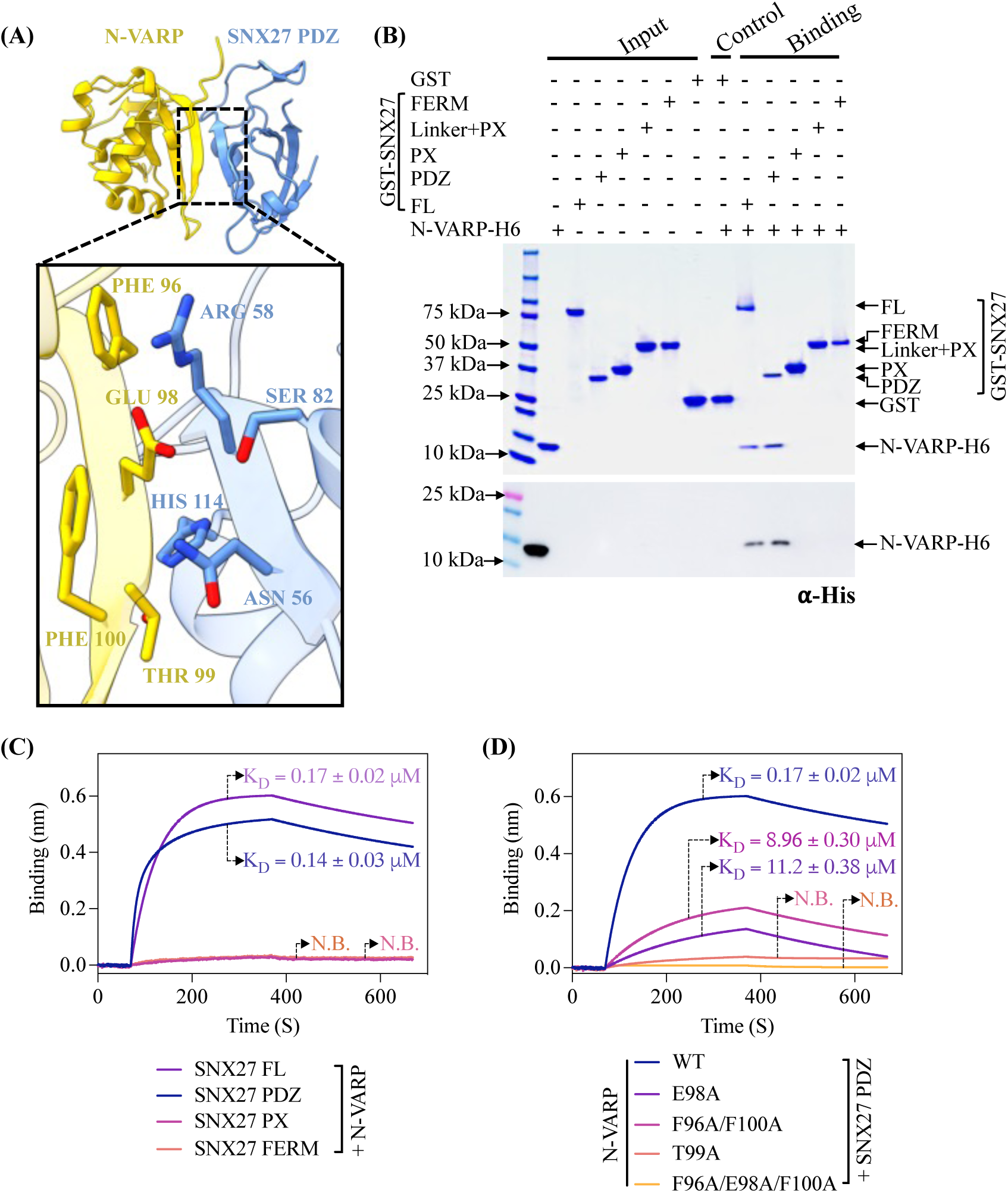
The VARP N-terminal domain directly binds SNX27 PDZ domain. **(A)** Ribbon diagrams of AlphaFold2.3 Multimer complex model between N-VARP (gold) and SNX27 PDZ (sky blue). The boxed inset shows interacting side chain residues on N-VARP (gold sticks) and SNX27 PDZ (blue sticks). **(B)** Pulldown experiments with purified GST-SNX27 fusion proteins. SNX27 full-length (FL), PDZ, PX, or FERM domains were used as bait with N-VARP-H6 as prey. A representative SDS-PAGE gel stained with Coomassie blue is shown in top panel with ⍺-His Western blot shown in bottom panel. **(C)** Biophysical data using BLI Octet system reveal a low micromolar affinity between SNX27 and N-VARP. Biotinylated N-VARP was loaded onto Streptavidin biosensor for measurements with SNX27 purified proteins. **(D)** Hydrophobic residues on VARP drive binding to SNX27. Biotinylated N-VARP mutants (E98A; T99A; F96A/F100A; F96A/E98A/F100A) were loaded onto Streptavidin biosensor for measurements with the SNX27 PDZ domain. Fitted data were plotted in GraphPad Prism, and binding kinetics were calculated using the Octet R8 analysis software package.

**Table 2.** Molprobity evaluation SNX27 PDZ domain models with binding partners. The experimental structure of 5-HT4(a)R PDZbm motif with SNX27 PDZ domain (PDB ID: 5EM9) is compared with the AlphaFold2.3 Multimer model of N-VARP bound to SNX27 PDZ domain reported here.

|  |  | PDB ID **5EM9**<br>**5-HT4(a)R:SNX27 PDZ** | AlphaFold2.3 Multimer<br>**N-VARP:SNX27 PDZ** |
| --- | --- | --- | --- |
| All atom contacts | Clashscore (all atoms) | 5.03 | 2.44 |
| Protein geometry | Poor rotamers | 0 (0%) | 2 (1.11%) |
|  | Favored rotamers | 82 (97.62%) | 176 (97.78%) |
|  | Ramachandran outliers | 0 (0%) | 0 (0%) |
|  | Ramachandran favored | 94 (96.91%) | 201 (97.10%) |
|  | Ramachandran distribution (Z-score) | 0.64 ± 0.85 | -0.12 ± 0.52 |
|  | MolProbity score<sup>a</sup> | 1.45 | 1.22 |
| | Cβ deviation >0.25 Å | 0 (0%) | 0 (0%) |
|  | Bad bonds | 0/792 (0%) | 0/1655 (0%) |
|  | Bad angles | 0/1074 (0%) | 4/2239 (0.18%) |
| Peptide omegas | Cis prolines | 0/3 (0%) | 0/9 (0%) |
| Additional validations | Chiral volume outliers | 0/125 | 0/259 |

In columns, raw count is listed first with percentage in parentheses.
<sup>a</sup> MolProbity score combines the clashscore, rotamer, and Ramachandran evaluations into a single score.

We next validated AF2.3 models using GST pulldown assays with recombinant purified proteins. Full-length GST-SNX27 or GST-tagged SNX27 domains (PDZ, PX, FERM) were used as bait and His-tagged N-VARP as prey. Consistent with AF2.3 predictions, N-VARP could pull down both full-length SNX27 and the SNX27 PDZ domain, but no detectable binding occurred with SNX27 PX and FERM domains alone acting as baits (Figure 2B). This interaction was quantified using BLI (Figure 2C). N-VARP binding to either full-length SNX27 or the PDZ domain exhibit very similar K_D_ values, further indicating the PDZ domain mediates the interaction.

Remarkably, all AF2.3 computational models suggest a short sequence within N-VARP (residues 95-101, sequence: LFEETFY) binds the conserved SNX27 PDZbm pocket (Figure S4B; S5A). Comparison of the N-terminal VARP sequence with well-established PDZbm motifs revealed similarities with classical type I PDZbm sequences (D/E^-3^−S/T^-2^−X^-1^−Φ^0^, where Φ represents any hydrophobic residue) (Figure S5B). Structural analysis of AF2.3 models reveal specific residues (Figure 2A) that mediate the molecular interaction between SNX27 PDZ and N-VARP. VARP Thr99 (equivalent to PDZbm −2 position) sits in close proximity to SNX27 His114 in the PDZ domain. VARP Glu98 (equivalent to PDZbm −3 position) is located such that it could form electrostatic and hydrogen bonds with SNX27 Arg58 and Asn56, as well as a hydrogen bond with SNX27 Ser80. VARP Phe96 (equivalent to PDZbm −5 position) and Phe100 (equivalent to PDZbm −1 position) residues sit adjacent to SNX27 Arg58 and Asn56 residues (Figures 2A; S5A).

Finally, we introduced structure-based point mutations into the VARP N-terminus to test the necessity of specific residues. We generated two single mutants (VARP T99A and VARP E98A); a double mutant (VARP F96A/F100A); and a triple mutant (VARP F96A/E98A/F100A) for binding studies with the SNX27 PDZ domain in GST pulldown assays (Figure S5C) and BLI experiments (Figure 2D). Both the VARP T99A mutant and the F96A/E98A/F100A triple mutant exhibited no detectable binding to SNX27, while the E98A and F96A/F100A mutants displayed reduced binding compared to the wild-type N-VARP (Figure 2D). This suggests VARP Thr99 plays a central role in formation of the VARP and SNX27 complex, while VARP residues Glu98, Phe96, and Phe100 play important auxiliary roles in establishing contacts with SNX27.

### SNX27/Retromer tubulates membranes enriched with PI(3)*P* lipid and PDZbm cargo

The next goal was to establish the role of VARP in the context of endosomal coat protein assembly on membranes to establish which combinations of Retromer, sorting nexins, and VARP can bind and tubulate membranes *in vitro.* Several combinations of SNX proteins with and without Retromer form tubules *in vitro*, including mammalian SNX1/SNX5 (*33*), yeast and mammalian SNX3/Retromer (*20*), and yeast Vps5/Retromer (*42*). We developed a biochemical reconstitution system using purified mammalian recombinant proteins and cargo motifs (CI-MPR or PDZbm) together with liposomes containing lipid headgroups that mimic physiological compositions (Folch I, PI(3)*P*). These components were used to conduct liposome pelleting assays (Figures 3, 4, S6) paired with negative stain electron microscopy to visualize and measure diameters of observed tubules.

**Figure 3.**
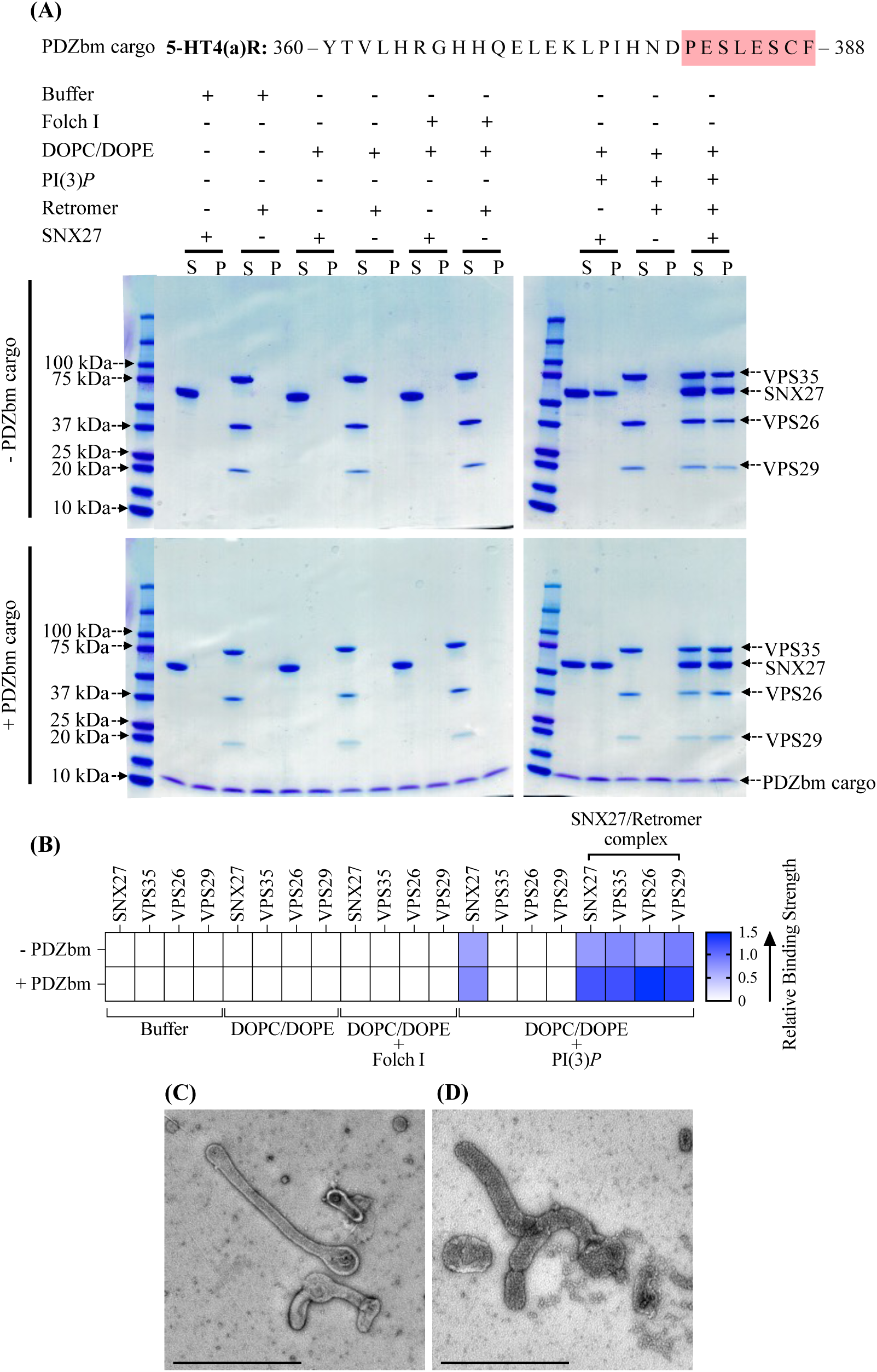
SNX27 and SNX27/Retromer can tubulate membranes in the presence of physiological lipid and cargo composition. (A) Liposome pelleting assays demonstrate membrane binding of human SNX27 alone and in the presence of Retromer. Purified recombinant SNX27, Retromer, and SNX27/Retromer complexes were incubated with or without liposomes enriched for PI(3)*P*, in the presence or absence of PDZbm cargo from the 5HT4(a)R family (residues 360–388; C-terminal PDZbm sequence highlighted in red). Buffer and DOPC/DOPE were used as negative controls to detect non-specific binding while Folch I was used to detect broad membrane-binding activity. Samples were subjected to ultracentrifugation followed by SDS-PAGE and Coomassie Blue staining of unbound supernatant (S) and bound pellet (P) fractions. (B) Protein complex binding to phosphoinositide-enriched membranes visualized by SDS-PAGE was quantified by measuring relative protein band intensities (ImageJ). The enrichment ratio between pellet and supernatant (P/S) was calculated for each protein band in the presence and absence of PDZbm cargo; relative intensity data are plotted as a heat map. (C) Imaging by negative stain EM reveals robust tubulation of PI(3)*P*-enriched liposomes incubated with SNX27 alone in presence of PDZbm cargo. (D) Negative stain EM indicates tubulation of PI(3)*P*-enriched liposomes incubated with SNX27/Retromer in the presence of PDZbm cargo motif from 5HT4(a)R. (Scale bars = 500 nm).

SNX27/Retromer is widely regarded as an endosomal protein coat based on multiple lines of evidence (*4*, *11*, *12*, *22*, *29*, *43*, *54*), but the ability of this complex to bind membranes and generate tubules has not been directly demonstrated. We first established whether SNX27/Retromer alone can function this way. We conducted liposome pelleting assays with the endosomal head group PI(3)*P* in the presence and absence of PDZ binding motif (PDZbm) cargo (Figure 3A). We generated a purified soluble His-tagged PDZbm cargo protein from the 5-HT4(a)R receptor family because its high affinity toward the SNX27 PDZ domain has been previously established (*50*). The data indicate SNX27 is specifically recruited to PI(3)*P*, but not Folch I, enriched liposomes (Figure 3A). Retromer requires recruitment by SNX27 to bind PI(3)*P* membranes (Figures 3A; 3B). Finally, the presence of PDZbm cargo positively impacts the membrane recruitment of both SNX27 and Retromer (Figure 3A; relative binding strength shown in Figure 3B). The partial enrichment in liposome pelleting (Figure 3B) signifies the influence and contribution of the PDZbm cargo, although lipid composition seems to drive membrane binding. As expected, none of the proteins showed binding under various control conditions, including buffer or DOPC/DOPE alone (Figures 3; S7).

We next used negative-stain electron microscopy (EM) to ascertain whether SNX27/Retromer induces curvature after membrane binding to generate tubules (Figure 3C; 3D). Both SNX27 alone (Figure 3C) and SNX27/Retromer (Figure 3D) can drive tubule formation from liposomes enriched with PI(3)*P* and PDZbm cargo, but the tubules have noticeably different diameters. SNX27 tubules exhibit an average diameter of 38.0 ± 5.0 nm (n=20 tubules) (Figure 3C; Table 3), while SNX27/Retromer coats produced wider tubules having an average diameter of 80 ± 6.0 nm (n=50 tubules) (Figure 3D; Table 3). The control experiment using PI(3)*P* liposomes alone did not induce membrane tubulation (Figure S7A). Together, these results indicate for the first time how SNX27, with and without Retromer, can induce membrane curvature and tubule formation *in vitro.* The data further highlight how Retromer contributes to membrane remodeling, since its presence induces tubules having a substantially wider diameter.

**Table 3.**
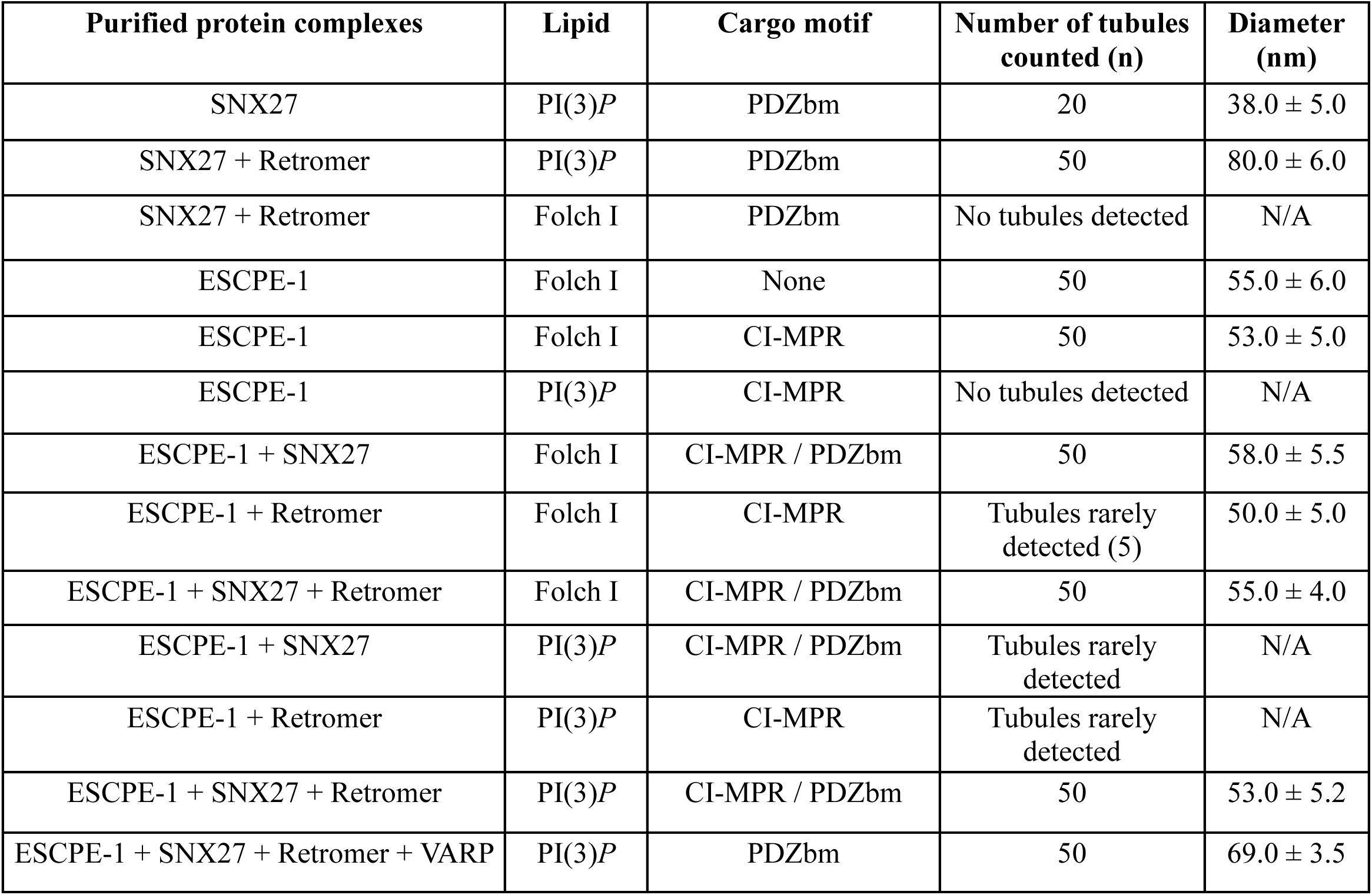
Measurements of membrane tubule diameters using negative stain Electron Microscopy (EM) in the presence of endosomal protein complexes in the presence of physiological lipid and cargo motifs.

### ESCPE-1 tubulates Folch I membranes displaying CI-MPR cargo

In endosomal coats, SNX-BAR heterodimers have been established as key players in deforming and tubulating membranes (*33*). We reconstituted the mammalian SNX2/SNX6 heterodimer, also known as ESCPE-1 (*33*), to ascertain whether and how it differs from SNX27/Retromer in its ability to bind and tubulate membranes with distinct compositions. ESCPE-1 specifically binds liposomes enriched with Folch I, emphasizing specificity for bis-phosphoinositides (PtdIns*P*_2_) over PI(3)*P* liposomes (Figures 4; S8A). SNX2/SNX6 exhibited no detectable pelleting in control buffer, DOPC/DOPE, or to PI(3)*P* membranes (Figure S8A; S8B). Negative stain EM revealed ESCPE-1 induces membrane tubulation of Folch I liposomes with average tubule diameters measuring 55.0 ± 6.0 nm (n = 50 tubules) (Figure S8C; Table 3). Control liposomes with Folch I alone did not induce tubulation (Figure S7B). Incorporating CI-MPR cargo into the SNX2/SNX6 and Folch I liposome mixture enhanced membrane binding (Figure S8A and S8B) and generated tubules with average diameter of 53.0 ± 5.0 nm (n = 50 tubules) (Figure S8D; Table 3). Notably, the tubule “run length” increased approximately 2-3 fold (Figure S8D) when CI-MPR cargo was present, highlighting its influence on ESCPE-1 tubule formation.

**Figure 4.**
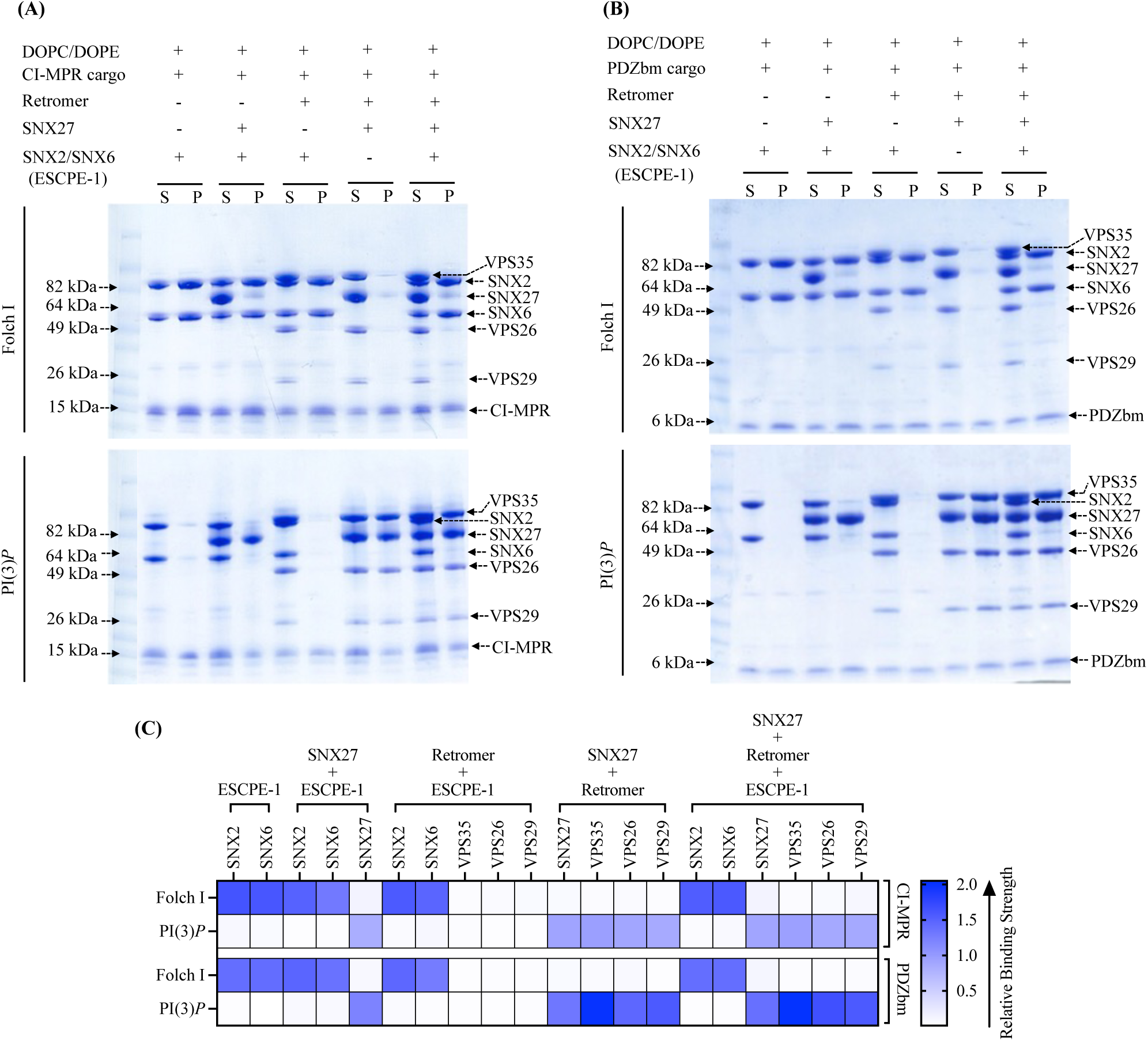
Biochemical reconstitution approaches reveal ESCPE-1 and SNX27/Retromer form different sub-complexes on membranes. Purified recombinant human SNX2/SNX6 (ESCPE-1), SNX27, and Retromer were incubated with **(A)** CI-MPR cargo motif or **(B)** PDZbm cargo motif from 5HT4(a)R in the presence of Folch I-(upper Coomassie gel) or PI(3)*P*-enriched liposomes (lower Coomassie gel). Samples were subjected to ultracentrifugation followed by SDS-PAGE and Coomassie staining of unbound supernatant (S) and bound pellet (P) fractions. **(C)** Protein binding to phosphoinositide-enriched membranes visualized by SDS-PAGE was quantified by measuring relative protein band intensities (ImageJ). Reconstitution data reveal specificity of sorting nexin complexes for both phospholipid and cargo composition on liposome membranes. SNX2/SNX6 (ESCPE-1) robustly binds membranes enriched in Folch I and CI-MPR cargo motifs, while SNX27 binds membranes loaded with PI(3)*P* and PDZbm cargo motifs. SNX27 recruits Retromer, while mammalian SNX2/SNX6 (ESCPE-1) complex does not appear to recruit Retromer in the presence of either cargo or phospholipid.

### Biochemical reconstitution approaches reveal ESCPE-1 and SNX27/Retromer form distinct subcomplexes having different tubule diameters

We next paired liposome pelleting assays with negative stain EM to assess which combinations of ESCPE-1, SNX27, and Retromer bind and tubulate membranes containing the phospholipid and cargo compositions established independently for SNX27/Retromer and ESCPE-1 (previous sections). One striking result was the inability of ESCPE-1 to recruit Retromer to Folch I membranes in presence of CI-MPR cargo; only SNX2/SNX6 was observed in the pellet fraction (Figure 4A, upper left Coomassie gel; Figure 4C, top row in heat map). We also tested Retromer recruitment in the presence of PDZbm cargo on Folch I membranes, since ESCPE-1 could presumably access this cargo in the context of an assembled supercomplex. However, Retromer is not observed in the pellet fraction (Figure 4B, upper right Coomassie gel; Figure 4C, third row in heat map). On PI(3)*P* liposomes, neither SNX2/SNX6 nor Retromer was detected in the pellet fraction (Figures 4A and 4B, lower Coomassie gel; Figures 4C, second and fourth row in heat map). Together, these data indicate the specificity of ESCPE-1 for Folch I membranes and support published models indicating ESCPE-1 alone may function as an independent coat.

Next, we assessed the influence of different sub-complexes on tubule morphology using negative stain EM. When Retromer was combined with SNX2/SNX6 in the presence of Folch I and CI-MPR cargo, there is a notable decrease in tubulation efficiency (Figure S9A, panel III), even though SNX2/SNX6 is robustly recruited to these membranes in pelleting assays (Figure 4A). SNX2/SNX6 alone consistently formed elongated tubules (Figure S9A, panel I), but adding Retromer with SNX2/SNX6 resulted only rarely in tubule formation with an average diameter of approximately 50 ± 5.0 nm (n = 5 tubules) (Figure S9A, panel III; Table 3). In line with liposome pelleting data, we rarely detect tubules with SNX2/SNX6 alone or in combination with Retromer on PI(3)*P* liposomes (Figure S9B, panels I and III; Table 3). Notably, control experiments revealed no detectable binding of Retromer alone to either Folch I or PI(3)*P* liposomes (Figure S7C). This striking negative result suggests mammalian SNX-BAR/Retromer system may have diverged away from its yeast counterpart.

Finally, we sought to establish whether ESCPE-1, SNX27, and Retromer forms an endosomal supercomplex in the presence of cargo and phospholipids. N-terminal extensions of SNX/BAR family members, including SNX1 and SNX2, have been shown to bind the SNX27 FERM module (Figure S1B, S1D) (*31*, *38*, *48*). However, we could not detect efficient pelleting of either SNX27 or SNX27/Retromer complex with ESCPE-1 on Folch I liposomes in the presence of either cargo (CI-MPR or PDZbm) (Figures 4A and 4B, upper Coomassie gels; Figures 4C, top and third row in heat map). PI(3)*P*-enriched membranes, in the presence of both cargoes, exhibited specificity for SNX27 alone and for SNX27/Retromer, with no detectable ESCPE-1 observed in pellet fraction (Figures 4A and 4B, lower Coomassie gel; Figures 4C, second and fourth row in heat map). In summary, these data reveal ESCPE-1 and SNX27/Retromer bind membranes with distinct compositions, with the phospholipid as a major driver of recruitment in this reconstituted *in vitro* system.

As before, we analyzed negative stain EM grids containing membrane-assembled complexes. These images reveal membranes exposed to ESCPE-1 and SNX27 generated membrane tubules with an approximate diameter of 58.0 ± 5.5 nm (n = 50 tubules) on Folch I (Figure S9A panel II; Table 3) membranes, while tubules were rarely detected with PI(3)*P* (Figure S9B, panel II; Table 3). Assemblies with ESCPE-1 and SNX27/Retromer complex exhibited tubules with an average diameter of 55 ± 4.0 nm (n = 50) for Folch I (Figure S9A, panel IV; Table 3) and 53 ± 5.2 nm (n = 50) for PI(3)*P* (Figure S9B, panel IV; Table 3). Notably, both varieties of tubules exhibited a close resemblance to those formed by ESCPE-1 alone (average diameter 55.0 ± 6.0 nm; Figure S9A panel I; Table 3). This similarity further suggests these tubules may be primarily decorated with the ESCPE-1 complex.

### VARP is required to reconstitute the proposed endosomal supercomplex on membranes

The finding that ESCPE-1 cannot recruit SNX27/Retromer to liposome membranes (Figure 4A, 4B) raises an important question. One likely explanation for failure to observe supercomplex formation is that a protein component is absent that allows endosomal sub-complexes to bind each other. The newly identified interaction presented here between the VARP N-terminus and SNX27 prompted us to test addition of VARP to the biochemical reconstitution. VARP is specifically implicated in SNX27/Retromer recycling to the plasma membrane (*61*), so we conducted pelleting assays with PI(3)*P*-enriched liposomes in the presence of PDZbm cargo. VARP addition yields an approximately stoichiometric complex between ESCPE-1 and Retromer in the pellet fraction on PI(3)*P*-enriched membranes (Figure 5A, far right lane; Figure 5B). SNX27 appears at slightly higher abundance within this complex, while VARP itself is sub-stoichiometric. These data agree with biophysical data (Figure 1C) and published data indicating one VARP binds two Retromer complexes via VPS29 subunits (*63*). We screened negative stain EM grids containing the full suite of endosomal proteins (SNX27, Retromer, ESCPE-1, and VARP) on liposomes with PI(3)*P* and PDZbm cargo. The observed tubules exhibit an average diameter of 69 ± 3.5 nm (n = 50 tubules) (Figure 5C), which is intermediate in size between tubules formed by SNX27/Retromer (Figure 3D) and ESCPE-1 complexes (Figure S9; Table 3). These data indicate how VARP incorporation induces changes in tubule diameter, which may arise from a change in coat lattice organization (see Discussion).

**Figure 5.**
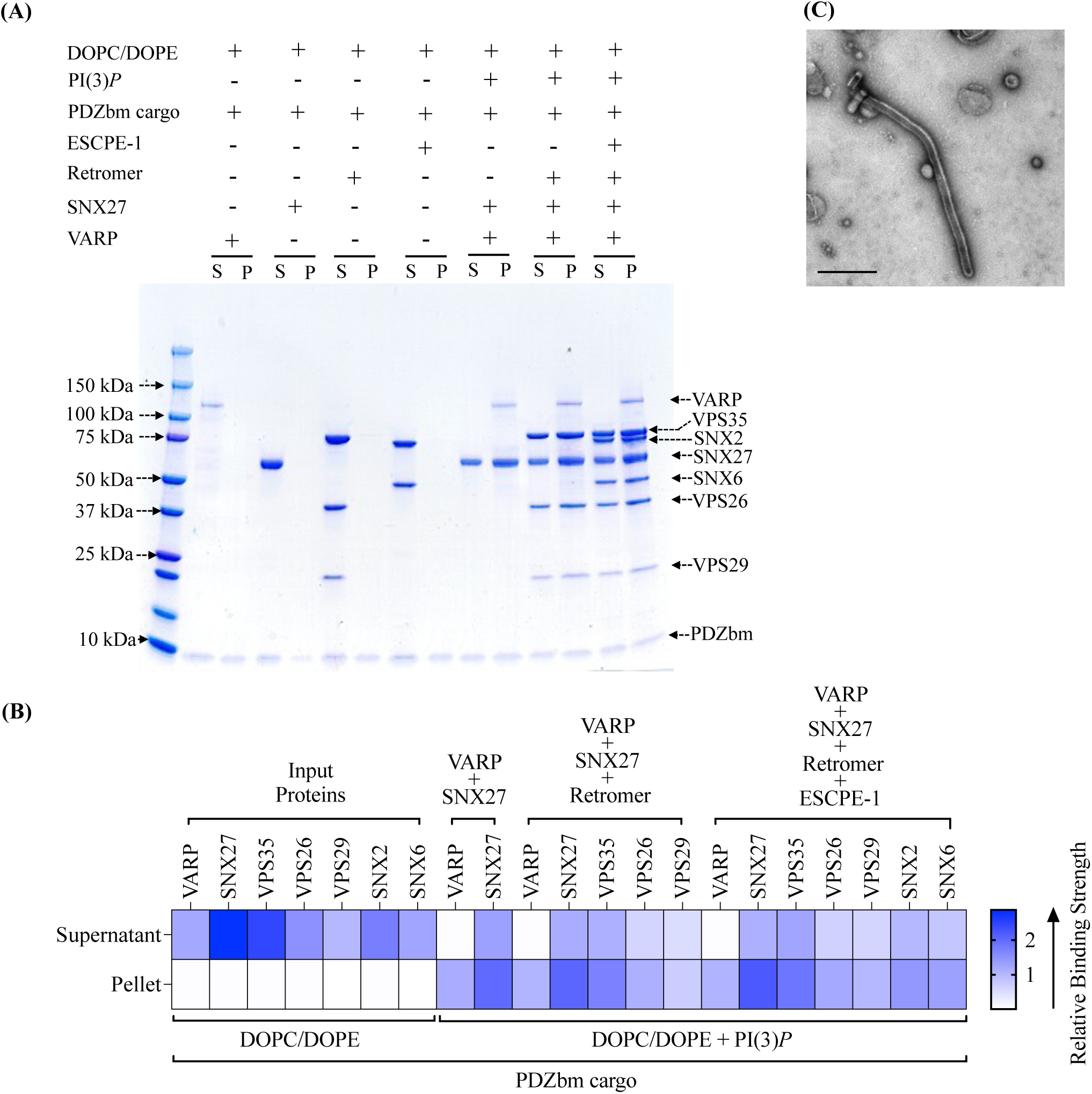
VARP is required *in vitro* to reconstitute the proposed SNX27/Retromer/ESCPE-1 endosomal ‘supercomplex’. **(A)** Purified recombinant human SNX27, Retromer, SNX2/SNX6 (ESCPE-1), and VARP were incubated with PDZbm cargo motif from 5HT4(a)R, either alone or together on PI(3)*P*-enriched liposomes. DOPC/DOPE was used as a negative control. Samples were subjected to ultracentrifugation followed by SDS-PAGE and Coomassie staining of unbound supernatant (S) and bound pellet (P) fractions. In the presence of VARP, all endosomal coat complexes (SNX2/SNX6, SNX27, and Retromer) are recruited to PI3*P*-enriched membranes. **(B)** Binding of proteins to phosphoinositide-enriched membranes visualized by SDS-PAGE was quantified by measuring relative protein band intensities (ImageJ). Relative gel band intensities corresponding to Supernatant (S) and Pellet (P) fractions were calculated for each protein sample and plotted as a heat map. **(C)** Representative negative stain EM image visualizing PI(3)*P* liposomes incubated with the SNX27/Retromer/ ESCPE-1/VARP ‘supercomplex’ in presence of the PDZbm cargo (scale bar = 500 nm). The supercomplex can both assemble **(A)** and tubulate membranes in the presence of VARP **(C)**.

Finally, we tested whether N-VARP alone is required to promote endosomal supercomplex assembly on membranes (Figure 6). Two approaches were undertaken to test this idea, because computational structural models and biophysical data suggest N-VARP would compete with PDZbm cargo motif binding. First, structure-based N-VARP single (T99A) and triple (F96A/E98A/F100A) mutants were introduced into liposome pelleting assays (Figure 6A) in the presence of PDZbm cargo and PI(3)*P.* When either VARP mutant is present, ESCPE-1 fails to bind membranes in a manner reminiscent of pelleting assays conducted without VARP (Figure 4). Neither mutant can reconstitute an approximately stoichiometric supercomplex observed in the presence of full-length VARP (Figure 5A). The second approach involved competition experiments. Isothermal titration calorimetry (ITC) experiments (Figure S10) further confirm N-VARP and PDZbm cargo motifs bind the same location on the SNX27 PDZ domain. PDZbm cargo motifs titrated into SNX27 PDZ alone give well-established low micromolar binding affinities (Figure S10). However, when purified N-VARP is added to SNX27 PDZ in the cell in a 1:1 ratio, the titrated PDZbm cargo peptide exhibits no detectable binding (Figure S10; Table 4). A conceptually similar experiment conducted in the liposome pelleting assay reveals N-VARP does not impede recruitment of the endosomal supercomplex to membranes enriched with PI(3)*P* and PDZbm cargo (Figure 6B). Two versions of the competition experiment were designed in the pelleting assay. The first version included a pre-incubation mixing step between SNX27 and N-VARP (Figure 6B-I). All components of the endosomal supercomplex were pelleted, but we observe an excess amount of N-VARP and reduced PDZbm cargo in pellet fractions. The second experiment (Figure 6B-II) included a pre-incubation step between SNX27 and PDZbm cargo. Here, all protein components in the endosomal supercomplex pelleted efficiently with roughly stoichiometric amounts of N-VARP and PDZbm cargo observed in pellet fractions. These data suggest VARP does not impede PDZbm cargo inclusion in the context of assembled coats and prompt many important hypotheses to test regarding the regulatory role of VARP on endosomal membranes (see Discussion).

**Figure 6.**
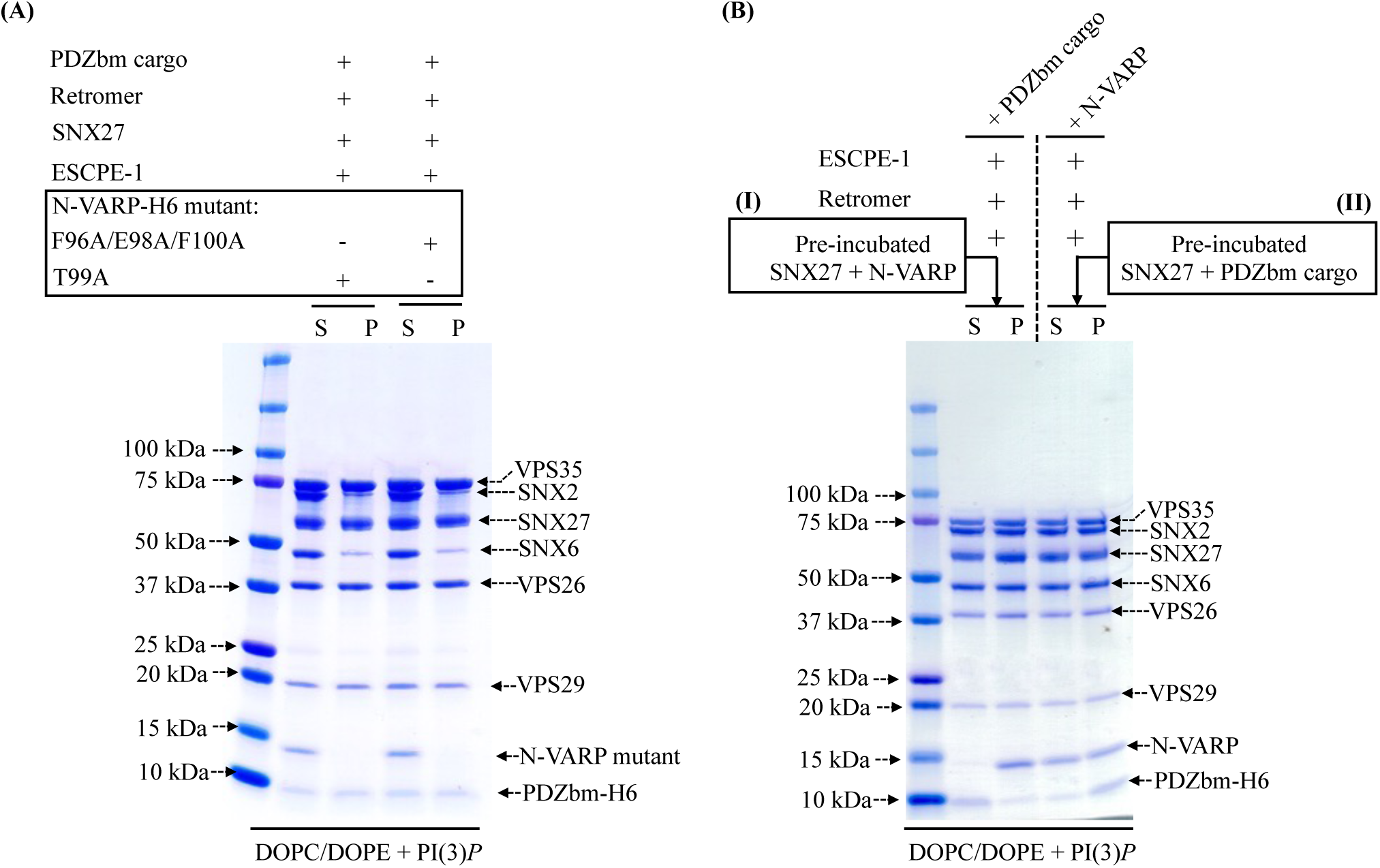
The VARP N-terminus is sufficient to recruit an endosomal supercomplex to membranes *in vitro.* **(A)** Liposome pelleting experiments demonstrate VARP N-terminus mutants (residues 1-117) cannot recruit the endosomal supercomplex to membranes *in vitro*. Purified proteins of the N-VARP triple mutant (F96A/E98A/F100A) or single mutant (T99A) were incubated with SNX27, ESCPE-1, and Retromer in the presence of PDZbm cargo and PI(3)*P*-enriched liposomes. In both experiments, SNX27 and Retromer are recruited, but ESCPE-1 exhibits only partial binding to membranes. N-VARP mutants remain in the supernatant (S) fraction. **(B)** A competition experiment demonstrates how binding between N-VARP and SNX27 does not interfere with PDZbm cargo binding and membrane recruitment in the liposome pelleting assays. In experiment (I), full-length purified SNX27 protein was pre-incubated (see Methods) with purified N-VARP protein. All endosomal coat protein components are pelleted efficiently in the presence of wild-type N-VARP. SNX27, Retromer, and ESCPE-1 are found in the pellet (P) fraction in a ration similar to that observed for full-length VARP (Figure 5). In experiment (II), full-length purified SNX27 protein was pre-incubated with purified PDZbm-H6 peptide. All other endosomal proteins were pelleted efficiently in the presence of N-VARP. In experiment (II), there is a greater amount of cargo in the pellet fraction compared to experiment (I). These results together suggest N-VARP is sufficient to promote endosomal supercomplex formation on membranes and does not inhibit cargo binding or incorporation into the coat.

**Table 4.** ITC binding data summary.

| Protein<br>(Cell) | Peptide<br>(Syringe) | K<sub>d</sub><br>(μM) | ΔH<br>(kJ/mol) | ΔS<br>(J/mol-K) | TΔS<br>(kJ/mol) | ΔG<br>(kJ/mol) | n |
| --- | --- | --- | --- | --- | --- | --- | --- |
| SNX27 PDZ<br>(25 μM) | 5-HT4(a)R-pS<sup>-5</sup> (500 μM) | 1.1 ± 0.2 | -63.6 ± 0.5 | - 99.2 ± 1.3 | - 29.5 ± 1.1 | - 34.6 ± 1.4 | 1.1 ± 0.1 |
| Pre-incubated<br>SNX27 PDZ<br>and N-VARP<br>(25 μM) | 5-HT4(a)R-pS<sup>-5</sup> (500 μM) | N.B. | N.B. | N.B. | N.B. | N.B. | N.B. |

## Discussion

### Summary

Cellular trafficking pathways rely heavily on interactions between and among multiple protein components to facilitate the sorting and transport of transmembrane protein cargo to their designated destinations. The endosomal system is particularly complex, with multiple protein players interacting across space and time to sort cargoes to different destinations. In this study, we aimed to elucidate how interactions between specific sorting nexin proteins (SNX2, SNX6, and SNX27) and the Retromer complex influence coat formation and membrane tubule morphology. Additionally, we established how VARP promotes formation of the previously proposed endosomal supercomplex composed of SNX27/Retromer and ESCPE-1 through a direct interaction with the SNX27 cargo binding PDZ domain. This biochemical reconstitution approach provides a powerful system to dissect other key protein-protein interactions and to provide testable hypotheses for cell-based experiments.

Published data have demonstrated VARP is recruited to endosomes through a direct interaction with Retromer (VPS29 subunit) and participates in the SNX27/Retromer recycling pathway that returns the glucose transporter GLUT1 to the plasma membrane (*54*, *61–65*). Work presented here provides biochemical evidence for an additional and new direct interaction between VARP and SNX27/Retromer coats through VARP binding to the SNX27 PDZ domain. Using pulldown assays and biolayer interferometry, we established VARP binds both Retromer and SNX27 with high nanomolar binding affinities. The stoichiometry between VARP and has been established previously (*63*) and here (as a positive control) as one VARP per two Retromers (Figure 1C). In contrast, BLI data clearly show VARP interacts with SNX27 with a 1:1 stoichiometry (Figure 1B).

### VARP N-terminus structure prediction and SNX27 binding

For the first time, we established a clear biochemical and biological role for the VARP N-terminus through direct binding to the SNX27 PDZ domain. Computational modeling using AF2.3 Multimer combined with with mutagenesis identified key residues on both VARP and the SNX27 PDZ that promote binding. Attempts to crystallize the VARP N-terminus have failed in our hands, but multiple versions of AlphaFold (data not shown) as well as models presented here (Figure 2, S2, S3) predict the N-terminus (residues 1-117) constitutes a small folded and globular domain. Notably, AF2.3 converges to a predicted model showing how N-VARP uses the sequence LFEETFY to bind the conserved SNX27 PDZ pocket known for its interaction with PDZ binding motifs in transmembrane receptors (*50*). Comparing the N-terminal VARP sequence with C-terminal PDZbm sequences from transmembrane receptors (Figure S5) reveals the VARP sequence resembles classical type I PDZbm motifs (D/E^-3^−S/T^-2^−X^-1^−Φ^0^; Φ represents any hydrophobic residue). The function of specific residues found in the VARP sequence would be consistent with prior data from experimental X-ray structures. For example, the Ser/Thr residue at the PDZbm −2 position is crucial for hydrogen bond formation with SNX27 PDZ residue His114 and essential for forming the complex between the SNX27 PDZ domain and PDZbm (*50*). Collaborative action of SNX27 residues Arg58, Asn56, and Ser80 provides a binding site for an acidic residue at the PDZbm −3 position. Accommodation of an acidic side chain adjacent to the −5 position by SNX27 Arg58 allows formation of an electrostatic plug. Structural analysis of the AF2.3 model between SNX27 PDZ and N-VARP reveals N-VARP residues likely maintain similar interactions with the SNX27 PDZ domain, including possible formation of specific hydrogen bonds and ion pairs. We leveraged AF2.3 models to guide mutagenesis studies (Figure 2D, 6A, S5C) and to independently validate the role of VARP residues in binding SNX27 (Figure 2, Figure S5). Overall, combining biochemical and biophysical approaches with computational modeling revealed a new molecular interaction in SNX27/Retromer coat complexes with important implications for coat assembly, cargo recognition, and regulation (discussed below).

### SNX27/Retromer assembly on membranes

A major goal for this study was to ascertain whether SNX27, alone and with Retromer, could remodel membranes to produce tubules in line with its proposed role as an endosomal coat. Using liposome pelleting assays and negative stain EM analysis, we demonstrated for the first time how SNX27 has inherent membrane-deforming capabilities and can induce tubulation of PI(3)*P*-enriched liposomes having relatively narrow average diameters (38 nm; summary in Figure 7). Addition of Retromer to the reconstitution results in coated tubules having significantly wider average diameters near 80 nm (Figure 7; Table 3). These differences highlight how SNX27 and Retromer interact cooperatively to remodel membranes and further support the idea that Retromer acts as a flexible scaffold (*19*). As expected, both phospholipid composition and cargo presence affect membrane recruitment of SNX27/Retromer. In the biochemical reconstitution system, phospholipid composition is a major driver for protein recruitment (Figures 3, 4), with cargo playing a role to enhance protein binding. SNX27 exhibits a clear “preference” for binding PI(3)*P* membranes over membranes containing bis-phosphoinositides in Folch I.

**Figure 7.**
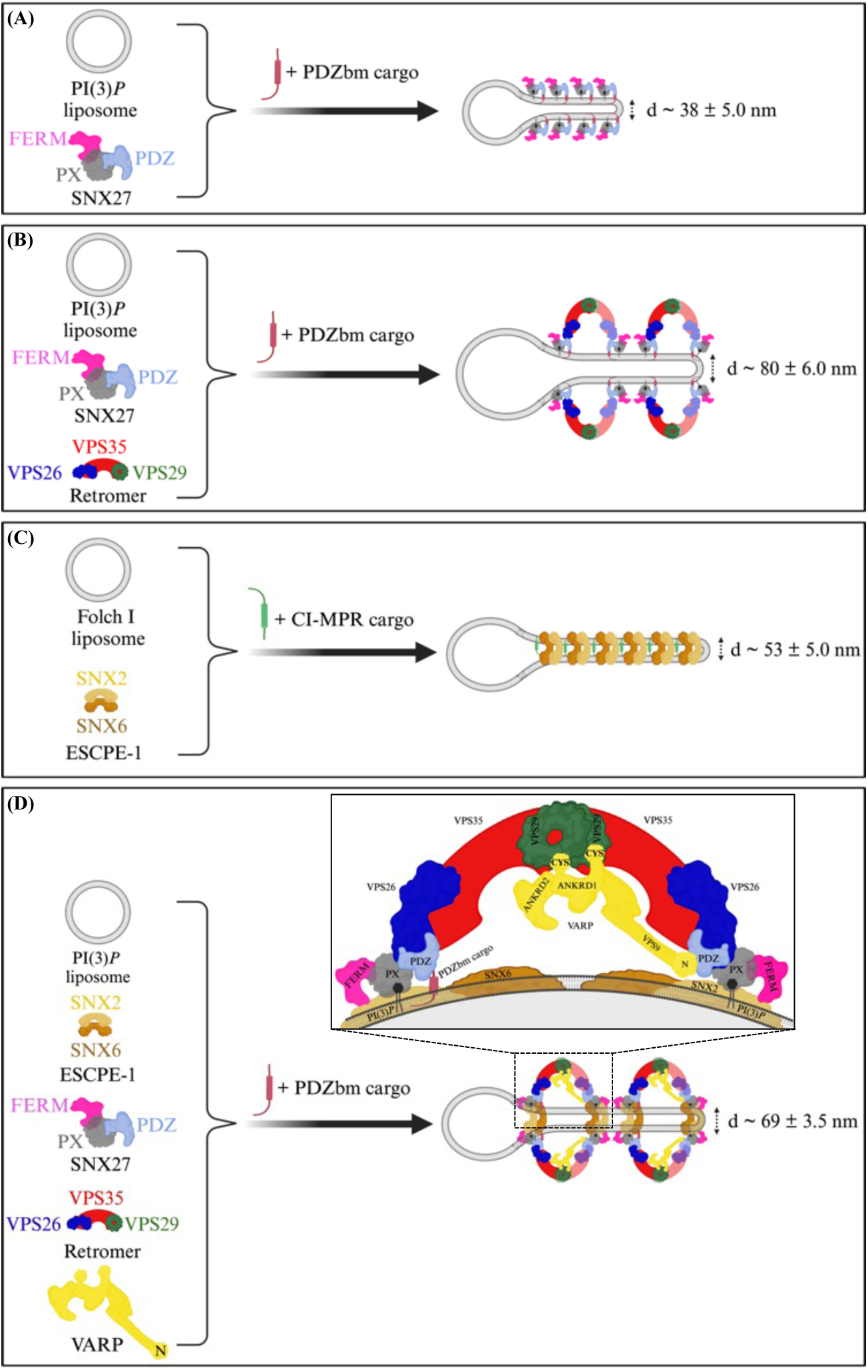
Morphologies of membrane tubules generated by endosomal coat complexes across different lipid and cargo compositions *in vitro*. Schematic comparison of endosomal coat complex combinations that generate tubules in the presence of physiological lipid and cargo compositions *in vitro*. Different endosomal coat proteins, either alone or as a complex, produce membrane tubules having different physical diameters as observed in negative stain EM (Table 3). SNX27 alone **(A)** or SNX27/Retromer **(B)** are shown here to generate tubules in the presence of PDZbm cargo and PI(3)*P*. **(C)** ESCPE-1 binds membrane containing Folch and CI-MPR cargo motifs, but does not interact with Retromer under these conditions. **(D)** The assembly of all individual coat components (SNX27, Retromer, and ESCPE-1) into the proposed endosomal ’supercomplex’ occurs on membranes with PI(3)*P* and PDZbm cargo motifs only in the presence of VARP. The stoichiometry quantified using BLI (Figure 1) suggests one VARP may bind one SNX27 and two Retromer copies in an arch and promote a conformation allowing the SNX27 FERM domain to bind the flexible SNX2 N-terminus. (Figure created using BioRender.)

The capability of SNX27 alone to deform membranes (Figure 3C) was unexpected. SNX27 lacks the canonical BAR domain that promotes dimerization and membrane binding in other SNX proteins, including SNX1, SNX2, SNX5, and SNX6. SNX3 also lacks a BAR domain and has been shown to bind Retromer and induce tubulation of PI(3)*P* membranes in the presence of Wntless (Wls) cargo peptide (*20*, *41*). Together, these results suggest Retromer-binding SNX proteins have multiple and different mechanisms to bind and shape endosomal membranes. SNX27 interacts with multiple protein and lipid membrane-associated ligands through its PDZ, PX, and FERM domains. It will be important to understand SNX27 architecture in the context of membrane binding to uncover the underlying mechanism that explains how it can remodel membranes containing PI(3)*P* specifically. In cells, SNX27 works together with Retromer to sort hundreds of important transmembrane receptors linked to neurological health (*12*, *50*, *54*, *55*), so understanding its role is important for both fundamental cell biology and human health.

### ESCPE-1 assembly on membranes

We employed ESCPE-1 in this study to establish how and when it can engage Retromer and SNX27 in a reconstitution system. These studies revealed a surprising negative result. For many years, the field has assumed the mammalian Retromer heterotrimer assembles under some circumstances with SNX1/SNX5 or SNX2/SNX6 to form a pentamer analogous to that observed in budding yeast. Mammalian SNX proteins arose from gene duplication and are orthologs to budding yeast Vps5 and Vps17. Recent data (*22*, *34*, *54*) suggest specific metazoan SNX-BAR proteins have diverged away from the Retromer heterotrimer, perhaps to form a separate coat called ESCPE-1. ESCPE-1 encompasses different combinations of SNX proteins (*5*, *29*, *31*), and CI-MPR is one important cargo. Here, we focused on SNX2/SNX6 as a model for ESCPE-1, because it was a tractable system for producing high quality purified proteins. The reconstitution data reproducibly demonstrate how ESCPE-1 robustly binds Folch I-enriched membranes but does not recruit Retromer. SNX27 and ESCPE-1 interact minimally in the absence of VARP, despite an established interaction between the SNX2 N-terminus and SNX27 FERM domain (*22*, *40*, *47*). However, we note there is a small interaction between SNX27 and ESCPE-1 observed in liposome pellet fractions (Figure 4), while there is essentially no observed Retromer binding. As with SNX27 (previous section), lipid composition is a major driver for ESCPE-1 membrane recruitment. ESCPE-1 robustly binds Folch I-enriched membranes, likely because the SNX2 PX domain exhibits stronger binding to bis-phosphoinositide headgroups including PI(3,4)*P*_2_ and PI(3,5)*P*_2_ (*26*, *69*, *70*). In contrast, the SNX1/SNX5 ESCPE-1 complex (*33*) has been shown to associate with PI(3)*P* but shows minimal interaction with PI(4,5)*P*_2_, PI(3,5)*P*_2_ and PI(3,4)*P*_2_. These data together suggest different ESCPE-1 complexes composed of distinct SNX heterodimers may recognize membranes having different phospholipid compositions; this could ensure cargo capture under dynamic lipid turnover conditions on endosomal membranes. Our results further demonstrate how ESCPE-1-decorated tubules reproducibly differ in diameter from SNX27 or SNX27/Retromer tubules (Table 3). ESCPE-1 forms tubules with average diameters near 53 nm (Figure 7) in the presence of CI-MPR cargo peptide. SNX27 cargo-loaded tubules are substantially more narrow (38 nm average diameter; Figure 7; Table 3) while SNX27/Retromer cargo-loaded tubules are much wider (80 nm average diameter; Figure 7; Table 3). CI-MPR is a cargo specific for ESCPE-1 through engaging SNX6 (*33–35*). As with SNX27/Retromer, incorporation of a cargo specific to ESCPE-1 somewhat enhances membrane binding and tubulation. The increased (2-3 fold) run-length of ESCPE-1 tubules in the presence of CI-MPR is especially striking (Figure S8D), although we were unable to robustly quantify this measurement.

### Implications for endosomal supercomplex assembly, regulation, and cargo sorting

A full biochemical reconstitution system allowed us to test an important unresolved question: can SNX27, Retromer, and ESCPE-1 form a proposed endosomal supercomplex (*18*, *21*)? ESCPE-1 alone is implicated in retrograde trafficking to the TGN, while SNX27/Retromer sorts cargoes to the plasma membrane. But ESCPE-1 has also been observed on recycling tubules (*12*), and reported interactions between the SNX1 and SNX2 flexible N-termini and SNX27 FERM domain further suggest direct binding (*22*, *40*, *47*). Liposome pelleting data revealed that lipid composition and cargo alone were insufficient to promote formation of the supercomplex (Figure 4). This result suggested two main possibilities: a supercomplex does not form, or a key regulatory component was missing.

VARP was a striking candidate as the missing component for several reasons. VARP binds Retromer (*61–63*); the R-SNARE, VAMP7 (*62*); and several Rab proteins (*61*, *64*). VARP uses its two small Cys-rich zinc motifs to bind VPS29, but its location within assembled coats is unclear. VARP has been implicated in GLUT1 recycling in SNX27/Retromer-mediated pathways (*54*). Addition of either full-length purified VARP or the N-terminus alone into the reconstitution revealed the first robust biochemical interaction among all components on membranes (Figure 5, 6) and suggests the endosomal supercomplex assembles under certain conditions. These data are insufficient for a robust quantitative approach to establish supercomplex stoichiometry, partly because some proteins (e.g. VPS29) stain less robustly than others. Nevertheless, the interaction among Retromer and ESCPE-1 appears roughly stoichiometric (Figure 5A, 5B), while there is approximately half as much VARP (Figure 5A, 5B). This is in line with biophysical data (Figure 1) and published work (*63*) suggesting one VARP engages two Retromers by interacting with VPS29 located at the top of assembled Retromer arches (*42*). In addition, there appears to be an excess of SNX27 relative to other components (Figure 5A, 5B). The assembled supercomplex reproducibly yielded tubules having average diameters of 69 nm (Table 3; Figure 7). The different size of these tubules from SNX27/Retromer tubules may suggest VARP regulates coat assembly by altering overall coat composition, architecture, or both. VARP may also introduce asymmetry into arches (discussed further below).

The data presented here prompt an important question regarding cargo recognition and binding in the context of assembled coats. The VARP N-terminus engages the well-established cargo binding pocket on the SNX27 PDZ domain (Figure 2, S4, S10) (*50*). This could suggest cargo and VARP compete for the same binding site on SNX27 PDZ. Calorimetry data here (Figure S10) and from other labs reveal low micromolar binding affinities (K_D_ 2-10 μM) and 1:1 stoichiometry between SNX27 PDZ domain and multiple PDZ binding motifs (*50*). BLI data (Figure 1B) reveal a 15-30x higher affinity for the interaction between SNX27 PDZ and VARP. In addition, the full supercomplex is robustly recruited to PI(3)*P* membranes that also contain PDZbm cargo in liposome pelleting assays (Figure 5, 6, S10). The data suggest VARP does not hinder cargo binding (at the very least). Another possibility is that VARP promotes a conformational change in SNX27 that is compatible with both PDZbm cargo inclusion and binding to SNX1 or SNX2 N-termini. In cells, there may be a multi-step process in which SNX27 is initially recruited to membranes with PI(3)*P* and PDZbm cargo. SNX27 is required to recruit Retromer (Figure 3), and both proteins harbor binding sites to recruit VARP. Rabs will also play an important role in establishing membrane identity and ensuring VARP recruitment. One critical role for VARP could be to ensure packaging of the R-SNARE, VAMP7, into the coat for a downstream fusion event. VARP may displace direct PDZbm cargo binding to SNX27 because N-VARP exhibits higher affinity for the PDZ domain. The stoichiometry of the VARP: SNX27 interaction is 1:1, suggesting VARP may displace only one PDZbm cargo (Figure 7) while simultaneously incorporating VAMP7, which could bring asymmetry into arches observed in structural studies (Figure 7D). In liposome assays (Figure 5, 6), the presence of full-length or N-terminal VARP alone allows ESCPE-1 to pellet, possibly because the SNX27 FERM adopts a conformation capable of engaging or perhaps releasing the flexible SNX2 N-terminus. VARP addition reproducibly produces tubules that differ in diameter from SNX27/Retromer tubules alone (Table 3), which further suggest a conformational change in coat architecture. Overall, the combination of a full biochemical reconstitution system with computational and quantitative biophysical methods provides a powerful suite of tools to test how and when multiple endosomal proteins collaborate to generate tubules *in vitro.* These data can be used for testing hypotheses for how cells can build and regulate tubular transport carriers for efficient cargo sorting out of endosomes. The role of flexible N-termini in regulating coat architecture is beginning to emerge and will have important implications for eukaryotic cell biology. These ongoing studies have critical implications for human health, since Retromer is considered a viable and important therapeutic target that engages different molecular interfaces when trafficking different cargo proteins.

## Materials and Methods

### Molecular biology and cloning

Mammalian Retromer constructs (VPS29, VPS35, VPS26 subunits) were generated in the labs of David Owen and Brett Collins and have been published previously (*13*, *71*). The original mammalian VARP (ANKRD27) construct was generated in the labs of Paul Luzio and David Owen (*61*, *62*). Human Sorting Nexin proteins (SNX27, SNX2 and SNX6), N-terminal VARP (N-VARP) and C-terminal cytosolic tail of PDZbm cargo 5-HT4(a)R (residues 360-388; sequence YTVLHRGHHQELEKLPIHNDPESLESCF) and CI-MPR cargo (residues 2347-2375; sequence SNVSYKYSKVNKEEETDENETEWLMEEIQ), optimized for *Escherichia coli* expression, were synthesized by the Genscript Corporation (USA). All the constructs, except full-length human VARP (VARP FL), were cloned either into the pET28A vector with an N-terminal 6X-His tag or into the pGEX-4T-2 vector with an N-terminal GST tag for expression and purification. Full-length human VARP (VARP FL) was cloned into mammalian expression vector pcDNA3.4+ with C-terminal 10xHis-tag.

### Site-Directed Mutagenesis

A PCR-based method using the Quikchange mutagenesis kit (NEB) was used to generate N-VARP mutants (E98A, T99A, F96A/F100A and F96A/E98A/F100A) using a plasmid encoding wild-type N-VARP as the template. A pair of oligonucleotide primers containing the desired mutation were used for the PCRs. The template plasmid DNA was linearized by DpnI digestion before transformation into *Escherichia coli* strain DH5α. Mutations were verified by DNA sequence analysis.

### Recombinant protein expression and purification

All the plasmids used in the current study were transformed into BL21(DE3)/pLysS E. coli cells (Promega) and expressed in LB 2xTY broth at 37°C until the A_600nm_ reached 0.8. Cultures were induced with 0.5 mM isopropyl 1-thio-β-D-galactopyranoside (IPTG) and allowed to grow at 20°C overnight, and cells were harvested by centrifugation at 6000 × g for 10 min, at 4°C. The cell pellet was resuspended in lysis buffer (20 mM Tris pH 8.0, 200 mM NaCl, 100 units DnaseI, and 2 mM β-mercaptoethanol). The cells were lysed by mechanical disruption at 30 kpsi using a cell disrupter (Constant Systems, UK). The lysate was clarified by centrifugation at 104,350 × g for 30 min at 4°C and the supernatant was loaded onto a column containing Ni-NTA metal affinity resins (Millipore Sigma, USA) for His-tagged proteins. The column was thoroughly washed with lysis buffer containing 100−500 mM salt. Finally, the protein of interest was eluted with a linear gradient of imidazole (from 100 to 250 mM) in 20 mM Tris pH 8.0 and 200 mM NaCl. Fractions containing the desired protein, as revealed by sodium dodecyl sulfate−polyacrylamide gel electrophoresis (SDS−PAGE), were pooled, and dialyzed against gel filtration buffer (20 mM Tris pH 8.0, 100 mM NaCl and 2 mM dithiothreitol (DTT)). For GST-tagged constructs, the supernatant from the clarified lysate was loaded onto a column containing Glutathione–Sepharose 4B resin (Cytiva, USA). Again, the column was thoroughly washed with lysis buffer containing 100−500 mM salt and subsequently the GST-tagged protein was either eluted in 20 mM Tris pH 8.0, 200 mM NaCl, and 20 mM reduced glutathione or the GST tag was cleaved overnight using thrombin (for pGEX4T2 vector containing constructs) at room temperature and GST free protein was eluted using buffer containing 20 mM Tris pH 8.0 and 200 mM NaCl. The eluted affinity purified proteins (His-tagged or GST-tagged or GST-cleaved) were finally subjected to size exclusion chromatography using a Superdex-200 16/600 HilLoad column, pre-equilibrated with 20 mM Tris pH 8.0, 100 mM NaCl and 2 mM DTT, attached to an ÄKTA Pure system (GE Healthcare, USA). Fractions containing pure protein, as revealed by SDS−PAGE, were pooled, and concentrated using appropriate cutoff concentrator (Centricon, Millipore Sigma, USA) and stored at −80°C.

Mammalian VARP was transiently expressed using the Expi293 Expression System (Thermo Fisher, Waltham, MA). Cells were grown in 250 mL flasks to a volume of 75 x 10^6^ cells per flask, then transfected with 1.0 μg plasmid DNA per mL of culture using the ExpiFectamine 293 kit (Thermo Fisher, Waltham, MA). Cells were harvested between 68 and 75 hours post transfection and frozen at -20°C until use. The frozen cell pellet was resuspended in lysis buffer (20 mM Tris pH 8.5, 500 mM NaCl, 100 units DnaseI, 4 mM MgCl_2_, and 2 mM β-mercaptoethanol). The cells were lysed using a dounce homogenizer and subsequently passed through an 18-gauge needle five times. The lysate was clarified by centrifugation at 104,350 × g for 30 min at 4°C and the supernatant was loaded onto a column containing Ni-NTA metal affinity resins (Millipore Sigma, USA). The subsequent purification steps were carried out analogous to the purification of a His-tagged protein, as described above.

### Phospholipids

DOPC (1,2-dioleoyl-sn-glycero-3-phosphocholine), DOPE (1,2-dioleoyl-sn-glycero-3-phosphoethanolamine), DOPS (1,2-dioleoyl-sn-glycero-3-phospho-L-serine), and DGS-Ni-NTA (1,2-dioleoyl-sn-glycero-3-[(N-(5-amino-1-carboxypentyl) iminodiacetic acid) succinyl]) nickel salt were purchased from Avanti Polar Lipids. PI(3)*P* (dipalmitoyl-phosphatidylinositol-3-phosphate) was purchased from Echelon Biosciences and Folch I (crude brain extract) was purchased from Sigma.

### Liposome preparation

All the phosphoinositides were protonated prior to usage. In brief, powdered lipids were resuspended in chloroform (CHCl_3_) and dried under argon. Dried lipids were then left in a desiccator for 1 h to remove any remaining moisture. Dried lipids were resuspended in a mixture of CHCl_3_:Methanol (MeOH):1 N hydrochloric acid in a 2:1:0.01 molar ratio, and lipids were dried once again and allowed to desiccate. Lipids were then resuspended in CHCl_3_:MeOH in a 3:1 ratio and dried once again under argon. Finally, dried lipids were resuspended in CHCl_3_ and stored at −20°C.

Folch I liposomes were formulated by mixing DOPC, DOPE, DOPS and DGS-Ni-NTA in a molar ratio of 42:42:10:3 with 1 mg/ml of Folch I. Similarly, liposomes containing PI(3)*P* were prepared by mixing DOPC, DOPE, DOPS and DGS-Ni-NTA in a molar ratio of 42:42:10:3 with 3 mole percent of PI(3)*P*. Both types of liposomes were prepared in a buffer containing 20 mM Hepes-KOH pH 7.5, 200 mM NaCl, and 1 mM Tris (2-carboxyethyl) phosphine at a final concentration of 1.0 mg/ml by performing 5 cycles of freeze-thaw steps followed by extrusion through a 0.4-μm polycarbonate filter. Control liposomes were prepared by combining DOPC and DOPE at a molar ration of 80:20.

### Liposome pelleting

For liposome pelleting experiments, 0.5 mg/ml of either Folch I liposome, or PI(3)*P* liposome, or control DOPC/DOPE liposome were used with the individual protein/protein complex sample (s) to a final volume of 100 μl. Following protein concentration were used for the liposome pelleting experiment: For the liposome pelleting experiments, 2.5 μM Retromer complex; 5 μM SNX2/SNX6 heterodimeric complex (2-fold); 10 μM SNX27 (4-fold); and 100 μM cargo adaptors (PDZbm or CI-MPR) (40-fold) were combined. The reaction mixture containing protein (s), and liposome, in presence or absence of cargo adaptors, and with or without cargo adaptors were left at room temperature (25°C) for almost 1 hrs to allow for protein–liposome interaction. After incubation, the solution was centrifuged at 50,000xg for 45 min. Supernatant and pellet fractions were separated and the pellet was resuspended in a buffer containing 20 mM Hepes-KOH pH 7.5, 200 mM NaCl, and 1 mM Tris (2-carboxyethyl) phosphine. Samples were then collected for analysis separated on a precast 4–12% Tris-glycine gel (BIO-RAD) and stained with Coomassie. The binding of the protein–phosphoinositide interactions within the SDS-PAGE has been further quantified by measuring the protein band intensities in ImageJ (http://rsbweb.nih.gov/ij/). The enrichment of the fraction of Pellet/Supernatant (P/S) was calculated for each protein band, both in presence and absence of relevant cargo, and plotted as a heat map.

### Negative Stain EM

For tubulation assays, 0.5 mg/mL liposomes [Folch I or PI(3)*P*] were incubated with either 5 μM of each individual protein (SNX27 or Retromer or SNX2/SNX6) in the presence of their respective cargoes (PDZbm or CI-MPR) or with protein combinations (2.5 μM Retromer, 5 μM SNX2/SNX6, 10 μM SNX27 along with 100 μM of respective cargoes) for 4 hrs at room temperature (25°C). 10 μl sample aliquots (protein with liposomes or liposomes alone) were adsorbed to glow-discharged 400-mesh carbon-coated copper grids (Electron Microscopy Sciences, EMS, USA) and stained with 0.75% uranyl formate and 1% uranyl acetate. The grids were examined on a Tecnai FEI Thermo Fisher Morgagni 100kV transmission electron microscope and images were recorded on a 1K X 1K AMT charge-coupled device camera. Tubule diameters were quantified in ImageJ analysis software (http://rsbweb.nih.gov/ij/) as an average of three measurements along the tubule.

### GST pull down assays

1 nmol of full-length GST-tagged SNX27 was mixed with 1 nmol of full-length His-tagged VARP for 1 hr at 4 °C. The Protein mixture was then centrifuged at high speed to remove any precipitated proteins. The supernatant was then added to pre-equilibrated (20 mM Tris pH 8.0, 200 mM NaCl, 1 mM DTT) Glutathione Sepharose resin and allowed to mix for an additional 30 min at 4 °C. Beads were washed five times in the above buffer supplemented with 0.5% Triton X100 (Sigma Aldrich). Bound proteins were analyzed by Western blots using mouse anti-His antibody (Abcam).

### Bio-layer interferometry (BLI)

The kinetics of the protein-protein interactions were determined using the bio-layer interferometry from the BLI system (Sartorius Octet BLI Discovery). Protein-protein interactions were observed by immobilizing 0.05 mg/ml of His-tagged VARP FL on a Ni-NTA biosensor or 0.05 mg/ml of biotinylated N-VARP on a streptavidin biosensor. After immobilization, the sensor was washed with buffer containing 10 mM Tris pH 8.0, 150 mM NaCl, and 0.1% BSA to prevent non-specific association. Increasing concentrations of SNX27 FL (0.5, 1.0, 2.0 and 4.0 μM) and Retromer (0.25, 0.5, 1.0 and 2.0 μM) were added to the Ni-NTA biosensor; whereas, increasing concentration of SNX27 FL, PDZ, PX and FERM domains (0.25, 0.5, 1.0 and 2.0 μM) were added to the streptavidin biosensor. The binding changes (nm) were measured in separate experiments performed in triplicate. Proteins were then allowed to disassociate from the probe in the same buffer. The data were processed and plotted using the Octet R8 analysis software package. Data from runs with full-length VARP and SNX27 proteins exhibited better fitting with 1:1 stoichiometric binding model (R^2^ = 0.99) compared to 1:2 binding model (R^2^ = 0.87). Similarly, data from runs with full-length VARP and Retromer heterotrimer exhibited better fitting with 1:2 stoichiometric binding model (R^2^ = 0.99) compared to 1:1 binding model (R^2^ = 0.92).

### Isothermal titration calorimetry (ITC)

ITC measurements were conducted on a Nano-ITC instrument (TA Instruments) in buffer consisting of 20 mM Hepes (pH 7.5), 100 mM NaCl and 2 mM DTT. PDZbm cargo peptide 5-HT4(a)R-pS^-5^ (Phosphorylated at Serine -5 position; commercially synthesized from Genscript) was dissolved in 20 mM Hepes (pH 7.5), 100 mM NaCl and 2 mM DTT for use in ITC binding experiments. In one experiment, 5-HT4(a)R-pS^-5^ peptide was titrated with purified SNX27 PDZ domain; in a second experiment, 5-HT4(a)R-pS^-5^ peptide was titrated with pre-incubated mixture of purified SNX27 PDZ and N-VARP domain proteins. In a typical experimental setup, the sample cell was filled with 300 μL of SNX27 PDZ domain protein or pre-incubated mixture of SNX27 PDZ and N-VARP proteins. The syringe contained a 50 μL solution of 5-HT4(a)R-pS^-5^ synthetic peptide (residues 1330-1336, EpSLESCF). All solutions were degassed prior to being loaded into the cell. Aliquots (2 μL) of 0.5 mM peptide solution from the syringe were injected into a 25 μM SNX27 PDZ or pre-incubated mixture of SNX27 PDZ and N-VARP domain protein solution at 25^◦^C with an interval gap of 3 minutes and the syringe rotating at 150 rpm to ensure proper mixing. Data were analyzed using Nanoanalyser software (TA Instruments) to extract the thermodynamic parameters, ΔH^◦^, *K*d (1/Ka), and stoichiometry (n). The dissociation constant (*K*d), enthalpy of binding (ΔH^◦^), and stoichiometry (n) were obtained after fitting the integrated and normalized data to a single-site binding model. The apparent binding free energy (ΔG^◦^) and entropy (ΔS^◦^) were calculated from the relationships ΔG^◦^ = RTln(*K*d) and ΔG^◦^ = ΔH^◦^– TΔS^◦^. All experiments were performed in triplicate to ensure reproducibility; standard deviations are reported from three runs (Table 4).

### AlphaFold Multimer computational modeling and validation

To generate predicted models of the SNX27 and VARP complex, we used the AlphaFold2 Multimer neural network-based structural prediction method (*72–74*). For complex modeling, the sequence of human full-length SNX27 (residues 1-541; Uniprot database Q96L92) was modeled with mammalian full-length VARP (residues 1-1050; Uniprot database Q96NW4) or N-terminal VARP alone (residues 1-117). AlphaFold version 2.3.2 (AF2.3.2) computations were executed using the resources of the Advanced Computer Center for Research and Education (ACCRE) at Vanderbilt. Structural alignments and images were generated with Pymol (Schrodinger, USA) or Chimera (*75*). In all AlphaFold2 Multimer predictions, we applied four criteria to evaluate model reliability (*72–74*): predicted local difference distance test (pLDDT) scores for local structure accuracy; interface predicted template modelling (ipTM) scores for the accuracy of the predicted relative positions of the subunits forming the protein-protein complex; predicted aligned error (PAE) scores for distance error between residues; and consistency among 5 top ranked models for prediction convergence as judged by structural superposition. In most cases, consistency of top 5 aligned models agreed with the pLDDT, ipTM and PAE criteria. The AlphaFold2 Multimer structural model was further validated using PISA (*67*) to evaluate buried surface area and MolProbity (*68*) to evaluate protein geometry and clashes (Table 2).

## Acknowledgements

We sincerely thank Scott Collier, Melissa Chambers, and Mariam Haider for their support at the Center for Structural Biology Cryo-EM facility (V-CEM). We also appreciate the Advanced Computing Center for Research and Education (ACCRE) facility at Vanderbilt University for providing access to run AlphaFold2 Multimer jobs. We thank the Biophysical Instrumentation Core Facility at Vanderbilt University for granting access to the Octet R8 Bio-layer Interferometer. M.C., A.K.K., and L.P.J. are supported by the National Institutes of Health Grant R35GM119525. M.C. was partly supported by a Pearson Fellowship from the Vanderbilt Department of Biochemistry. M.G.J.F. was supported by the National Institutes of Health Grant R35GM139546. V-CEM is supported by an NIH S10 award OD030292-01 at Vanderbilt University.

## Author contributions

MC: Investigation, Methodology, Visualization, Writing – original draft, Writing – review & editing. AKK: Methodology, Writing – review & editing. MGHF: Investigation, Methodology, Writing – review & editing, Funding acquisition. Lauren P. Jackson: Conceptualization, Writing – original draft, Writing – review & editing, Supervision, Funding acquisition.

---

**Figure S1.**
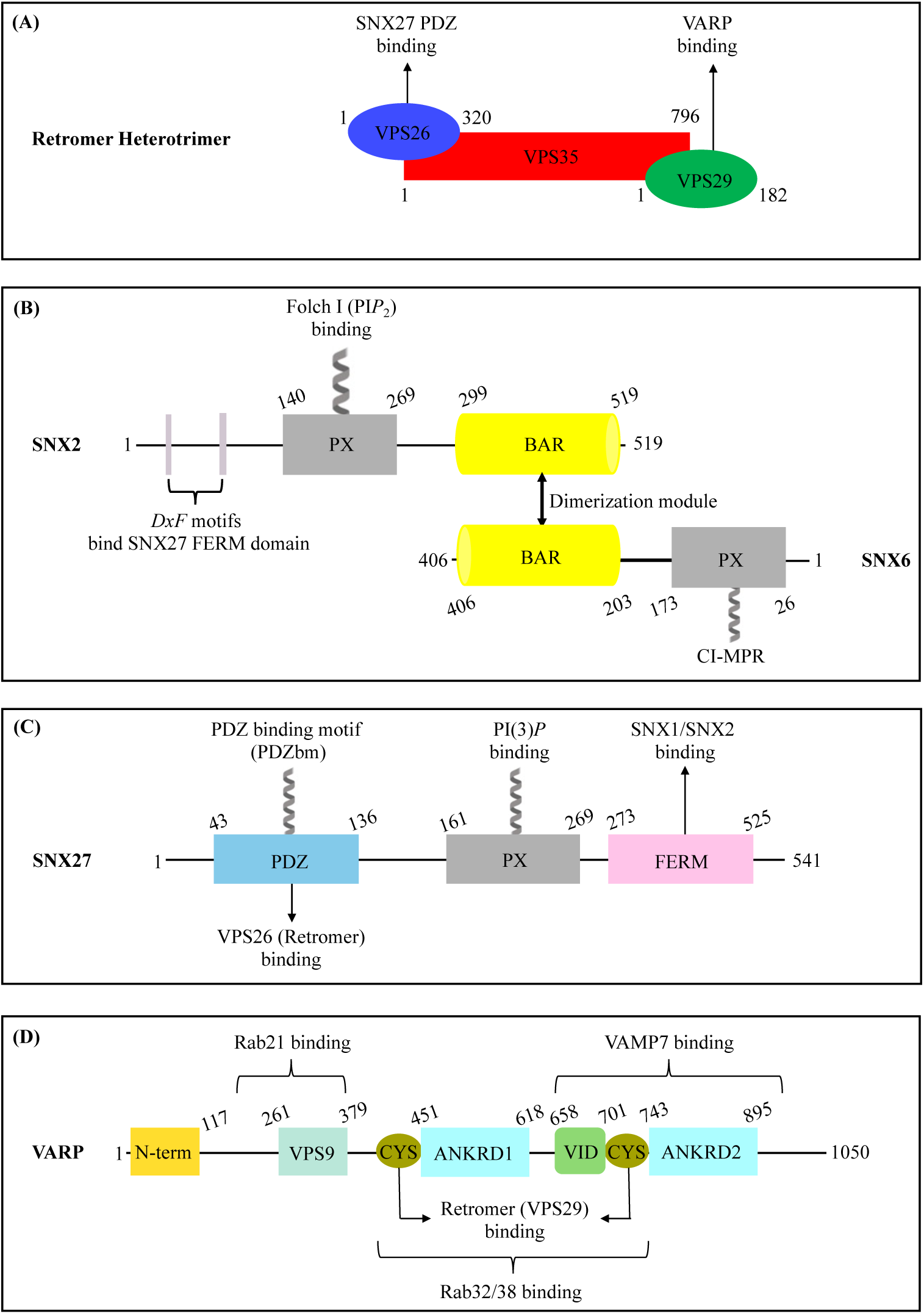
Domain architecture of endosomal proteins used in this study. **(A)** Domain architecture of mammalian Retromer containing three Vacuolar Protein Sorting (VPS) subunits: VPS26, VPS35, and VPS29. **(B)** Domain architecture of mammalian SNX2/SNX6 (ESCPE-1). The SNX2 N-terminus is an extended flexible region; the PX domain binds phospholipids; and the C-terminal BAR domain forms a dimer with SNX6. SNX6 contains a short and flexible N-terminus and a PX domain known to recognize a motif in CI-MPR cargo; and a C-terminal BAR domain. **(C)** Human SNX27 contains an N-terminal PDZ; central PX; and C-terminal FERM domains. The PDZ domain binds transmembrane proteins containing a PDZ binding motif (PDZbm) and the VPS26 subunit of Retromer. The PX domain recognizes PI(3)*P*, a the FERM module has been shown to bind flexible N-terminal regions of two SNX-BAR proteins, SNX1/SNX2. **(D)** Human VARP (also known as ANKRD27) domains are shown with established binding partners highlighted. Two cysteine-rich motifs (CYS) engage the VPS29 Retromer subunit, and this study establishes how the VARP N-terminus binds SNX27.

**Figure S2.**
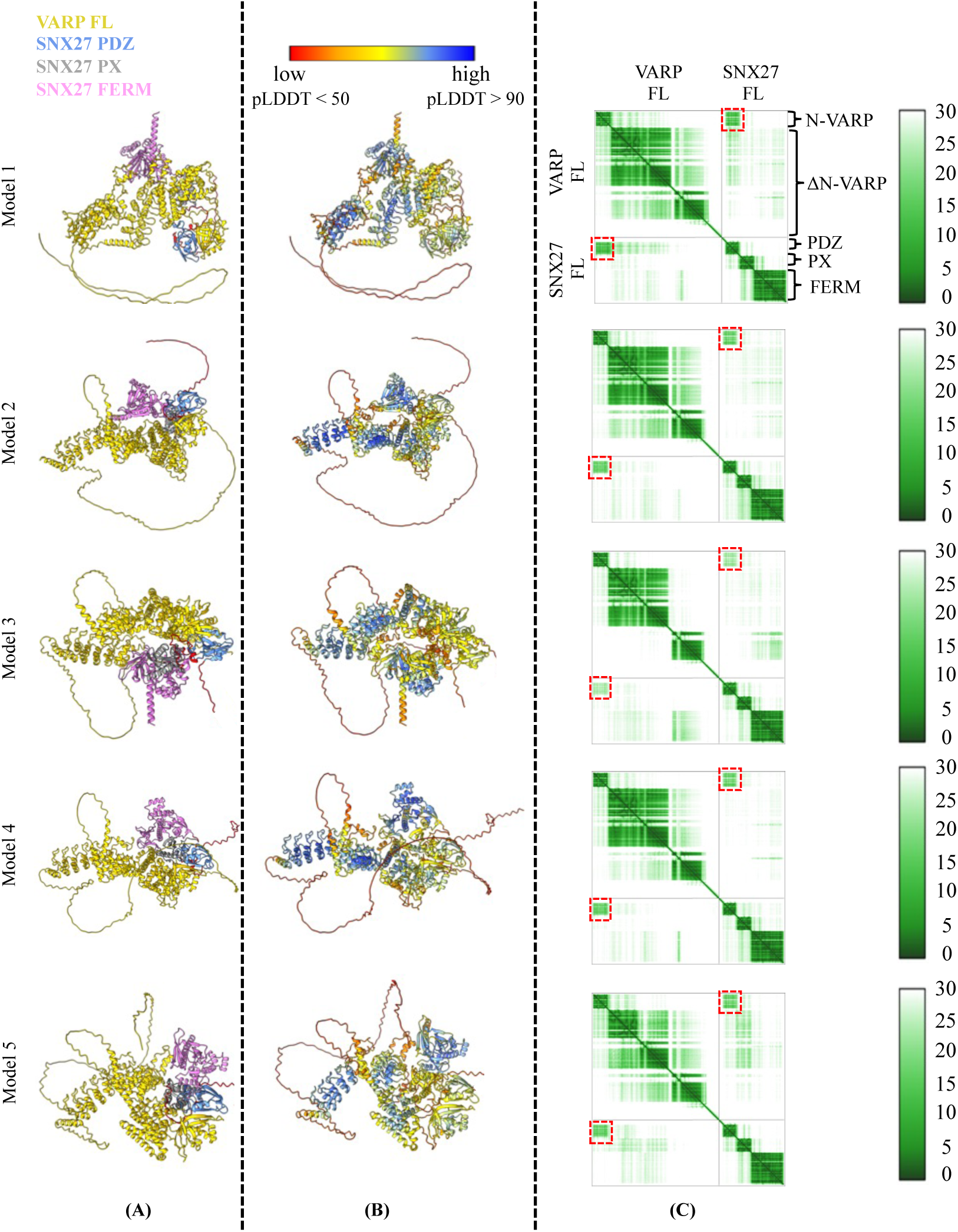
AlphaFold models of full-length VARP and full length SNX27. **(A)** Ribbon diagrams of the five top ranked models generated in AlphaFold2.3 Multimer depicting full-length VARP bound to full-length SNX27. VARP is shown in gold ribbons with SNX27 colored by domain: SNX27 PDZ in sky blue; SNX27 PX in grey, and SNX27 FERM in magenta color. **(B)** Ribbon diagrams of the top five models colored by pLDDT score. High pLDDT scores (shown in blue) reflect high confidence in local structure prediction. **(C)** For each model, the Predicted Aligned Error (PAE) score matrix is shown. Low scores (dark green color) represent high confidence in the relative position in 3D space (right column). The boundaries of protein domains can be observed in the PAE plots, including N-VARP (residues 1-117); ΔN-VARP (residues 118-1050); SNX27 PDZ (residues 43-136); SNX27 PX (residues 161-269); SNX27 FERM (residues 273-525). The predicted interaction between N-VARP and the SNX27 PDZ domain is highlighted as red dashed boxes on PAE plots.

**Figure S3.**
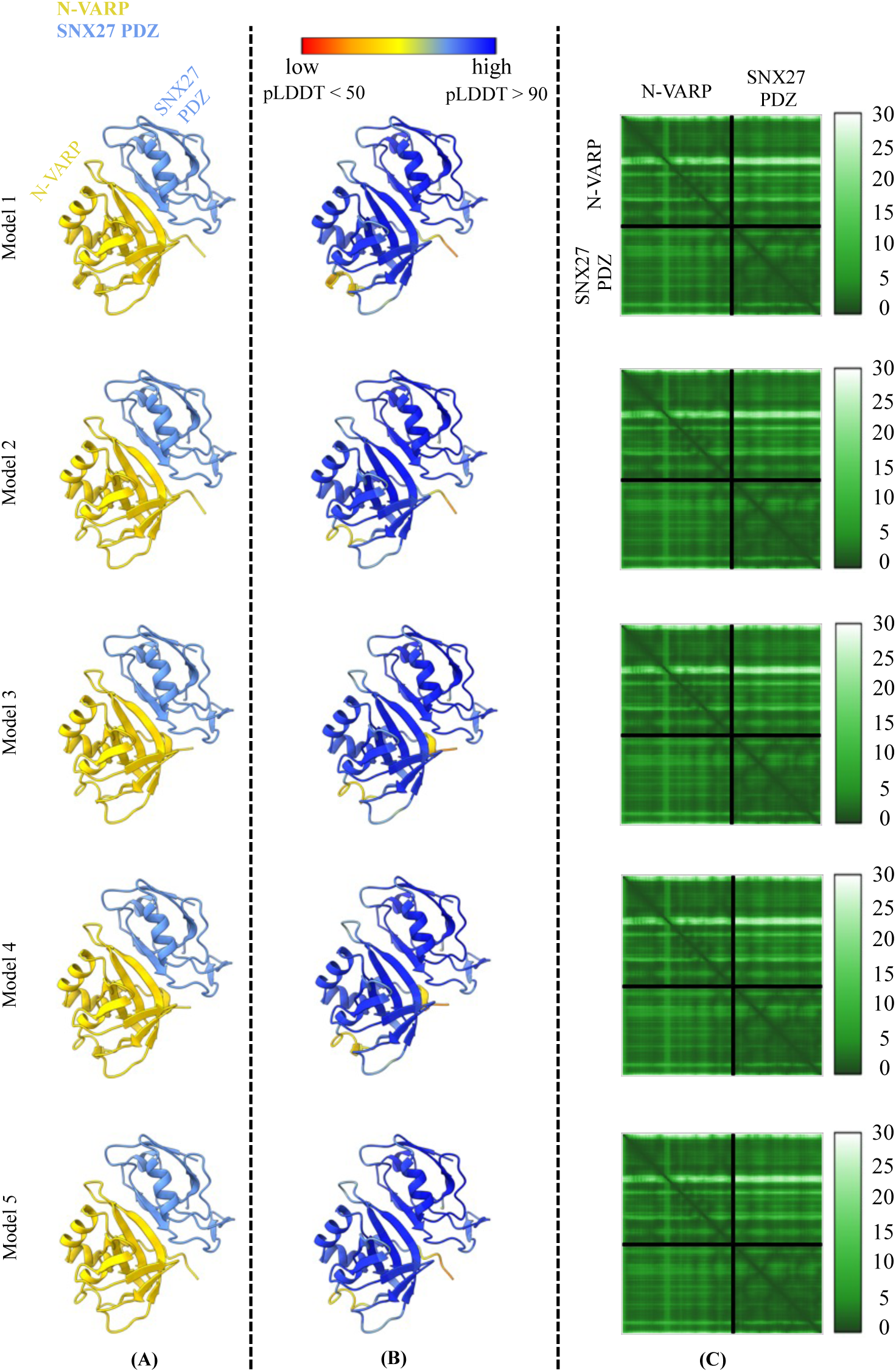
AlphaFold models of the VARP N-terminal globular domain with SNX27 PDZ domain. (A) Ribbon diagrams of the five top-ranked AlphaFold2.3 Multimer models depicting N-VARP bound to SNX27 PDZ. Models are colored by domain, with N-VARP in gold and SNX27 PDZ in sky blue. (B) Ribbon diagrams of top five models colored by pLDDT score; dark blue (scores >90) represents high confidence in local prediction. (C) For each model, the Predicted Aligned Error (PAE) score matrix is shown. The PAE score matrix provides low scores (deep green), signifying high confidence in the relative position in 3D space. The boundaries of protein domains (N-VARP and SNX27 PDZ) are labeled on PAE plots. Overall, AlphaFold consistently generates the same predicted model for this interaction.

**Figure S4.**
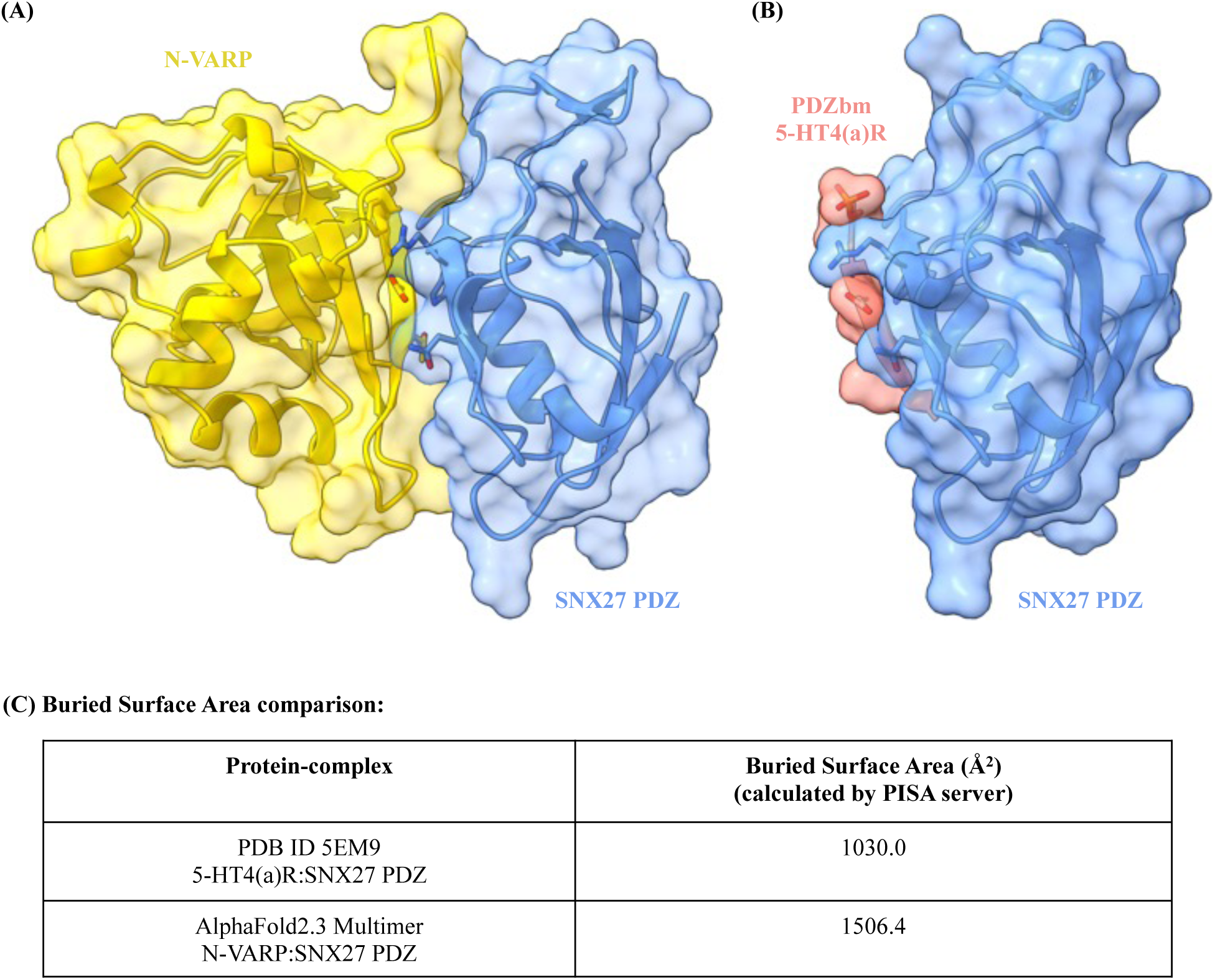
Comparative analysis of interactions between SNX27 PDZ and the VARP N-terminus or PDZ binding motif (PDZbm) cargo peptide. **(A)** Transparent surface view is shown over a ribbon diagram of VARP N-terminus (gold) and SNX27 PDZ domain (sky blue) model from AlphaFold. **(B)** Equivalent transparent surface view is shown over a ribbon diagram of the experimental X-ray structure with PDZbm peptide from 5-HT4(a)R (light red) bound to SNX27 PDZ domain (sky blue). (C) Comparison of predicted buried surface area of each structural model calculated in PISA. The interaction between N-VARP and SNX27 PDZ buries 50% greater surface area than does the PDZbm cargo peptide, in agreement with observed dissociation constants.

**Figure S5.**
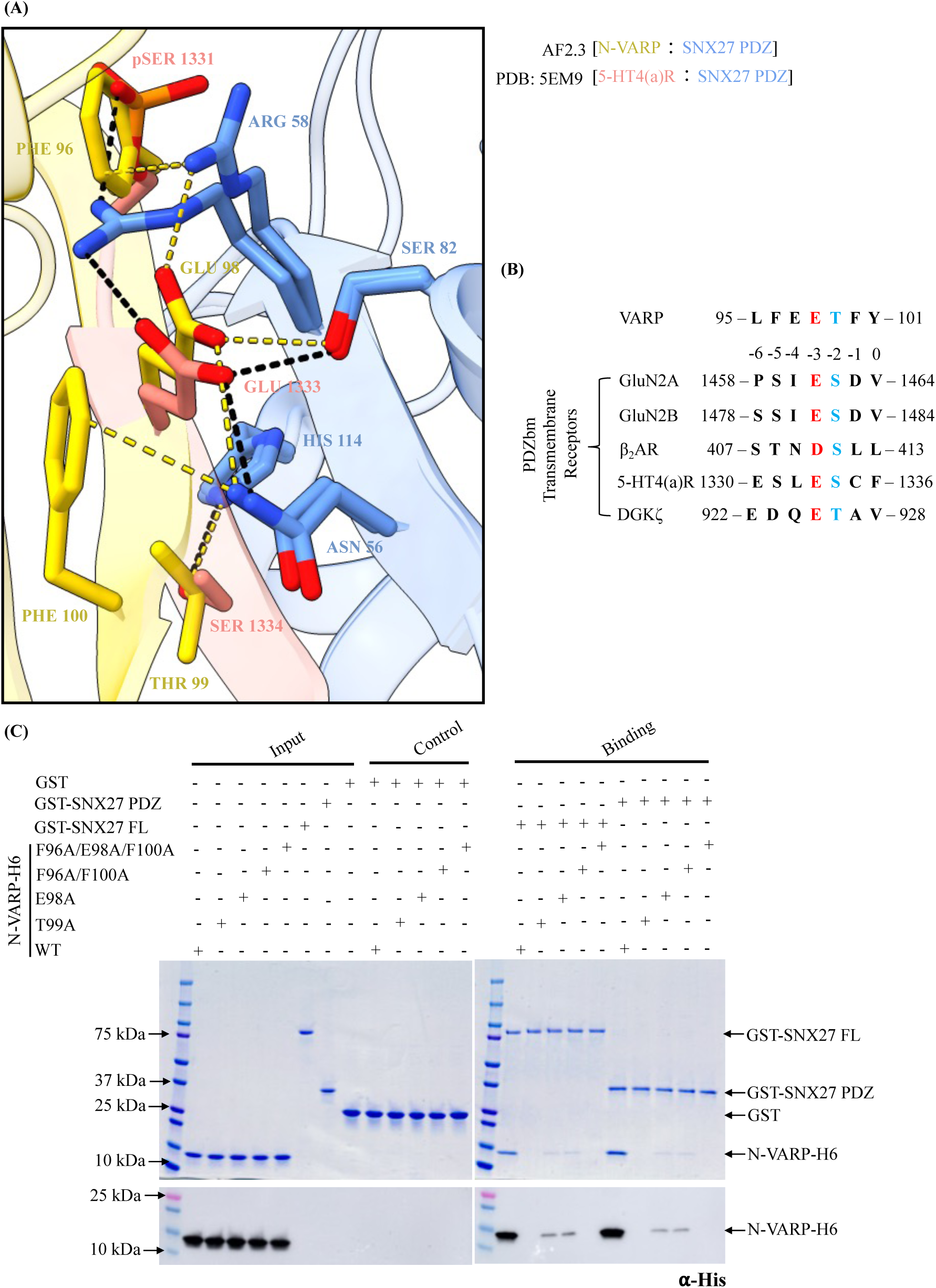
The VARP N-terminus and PDZbm cargoes bind in the same pocket on the SNX27 PDZ domain. (A) Close-up view of structural superposition between C-terminal PDZbm cargo from 5-HT4(a)R (light red side chains) and SNX27 PDZ domain (sky blue; PDB: 5EM9) and the AlphaFold-predicted model of N-terminal VARP (gold) bound to SNX27 PDZ (sky blue). The superposition reveals multiple overlapping residues that mediate both interactions. Both N-terminal VARP (residues Phe96, Glu98, Thr99 and Phe100) and phosphorylated PDZbm motif (residues phospho-Ser1331, Gle1333 and Ser1334) interact with the same patch on the SNX27 PDZ domain composed of residues Asn56, Arg58, Ser82, and His114. Predicted interaction distances between the VARP N-terminus and SNX27 PDZ domain are represented as yellow dashed lines, while distances determined from the experimental structure of the PDZbm cargo motif and SNX27 PDZ domain (PDB ID: 5EM9) are shown as black dashed lines. (B) Sequence alignment and comparison of VARP N-terminus (motif: LFEETFY; residues 95−101) and multiple PDZ binding motifs in five known transmembrane receptors. The motif position numbers are assigned according to the classical type I PDZbm sequence (D/E^-3^−S/T^-2^−X^-1^−Φ^0^, where Φ represents any hydrophobic residue). Residues corresponding to -2 and -3 positions are highlighted in blue and red, respectively. (C) GST pulldown experiments confirm VARP N-terminal residues from AlphaFold2 model are involved in binding. GST-tagged full length SNX27 or GST-SNX27 PDZ domain were used as baits with purified His-tagged N-terminal VARP mutant proteins (E98A; T99A; F96A/F100A double mutant; and F96A/E98A/F100A triple mutant) as prey. Representative SDS-PAGE gel stained with Coomassie blue is shown in top panel with ⍺-His Western blot shown in bottom panel.

**Figure S6.**
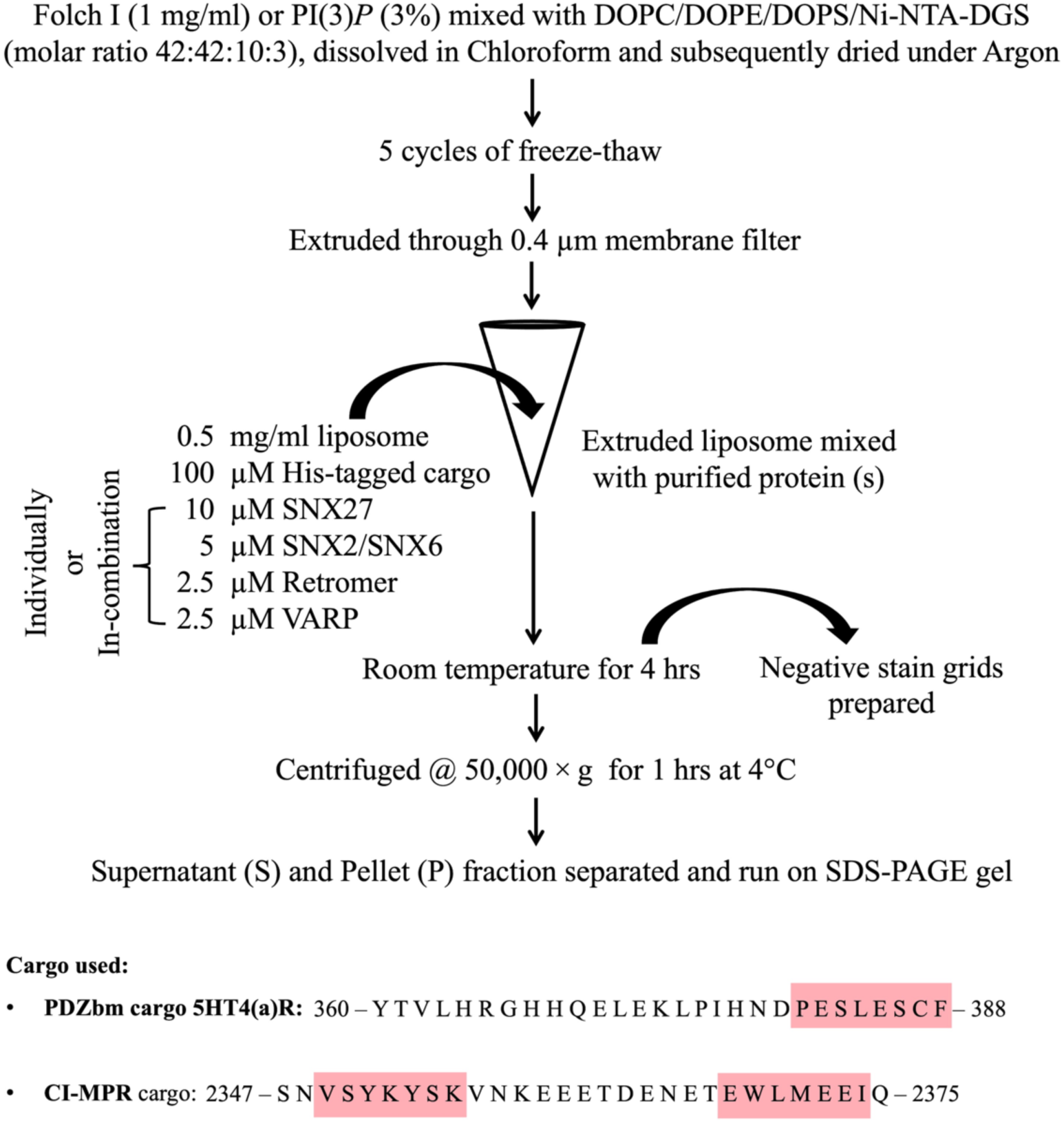
Flow chart depicting steps in liposome preparation and liposome pelleting assay.

**Figure S7.**
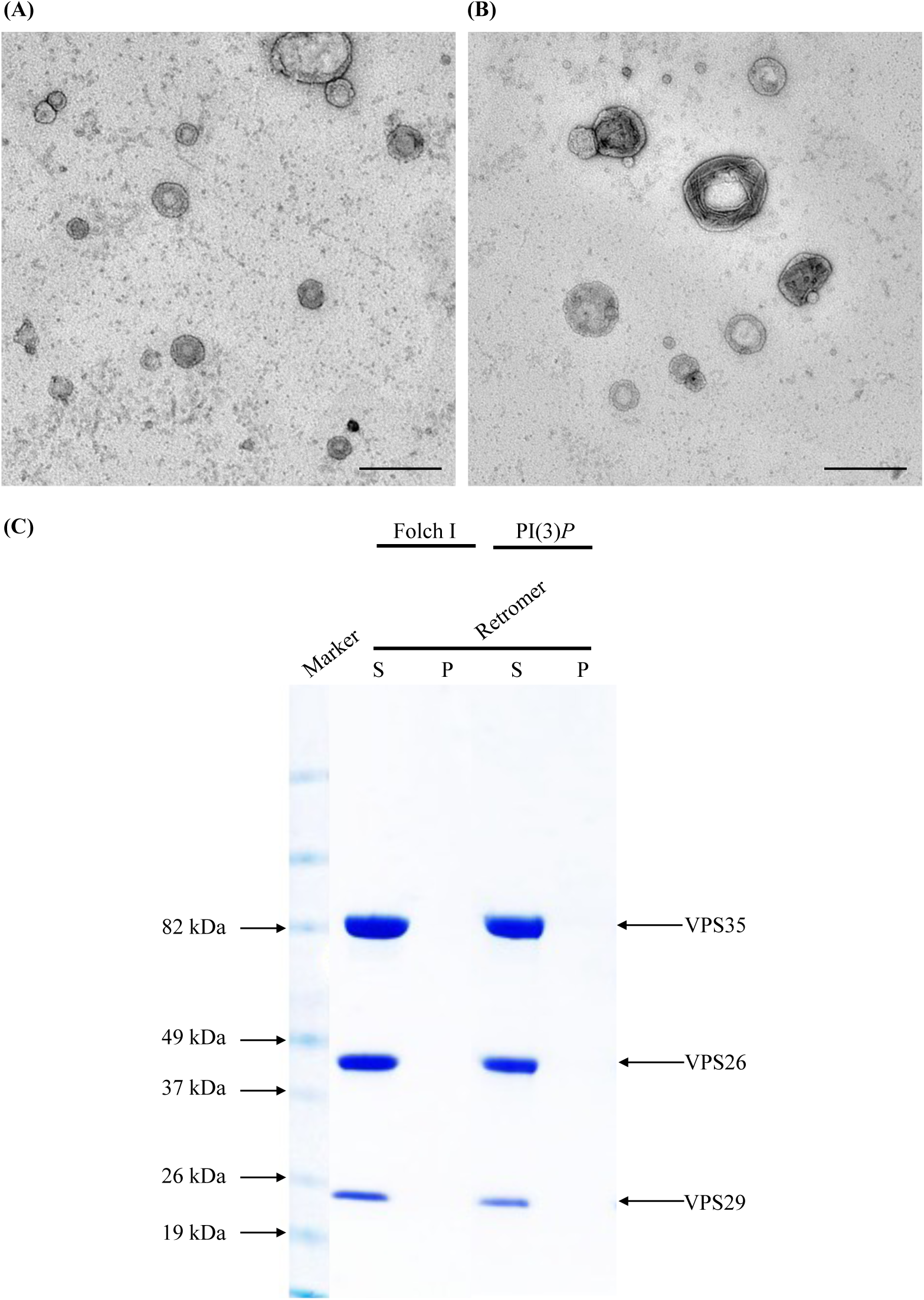
Control experiments to establish the reconstitution system. Representative negative stain EM images of liposomes containing (A) PI(3)*P* and (B) Folch I as controls. Liposomes containing these phospholipid compositions do not exhibit tubules. Liposomes containing either (A) PI(3)*P* or (B) Folch I were incubated with buffer (20 mM HEPES-KOH pH 7.5, 200 mM NaCl, and 1 mM Tris (2-carboxyethyl)phosphine) and visualized using negative stain EM. (Scale bar = 500 nm). (C) Liposome pelleting assay of purified Retromer complex on liposomes enriched with with Folch I or PI(3)*P*. Samples were subjected to ultracentrifugation followed by SDS-PAGE and Coomassie staining of the unbound supernatant (S) and bound pellet (P) fractions. Retromer is not recruited to membranes in the absence of cargo or SNX27.

**Figure S8.**
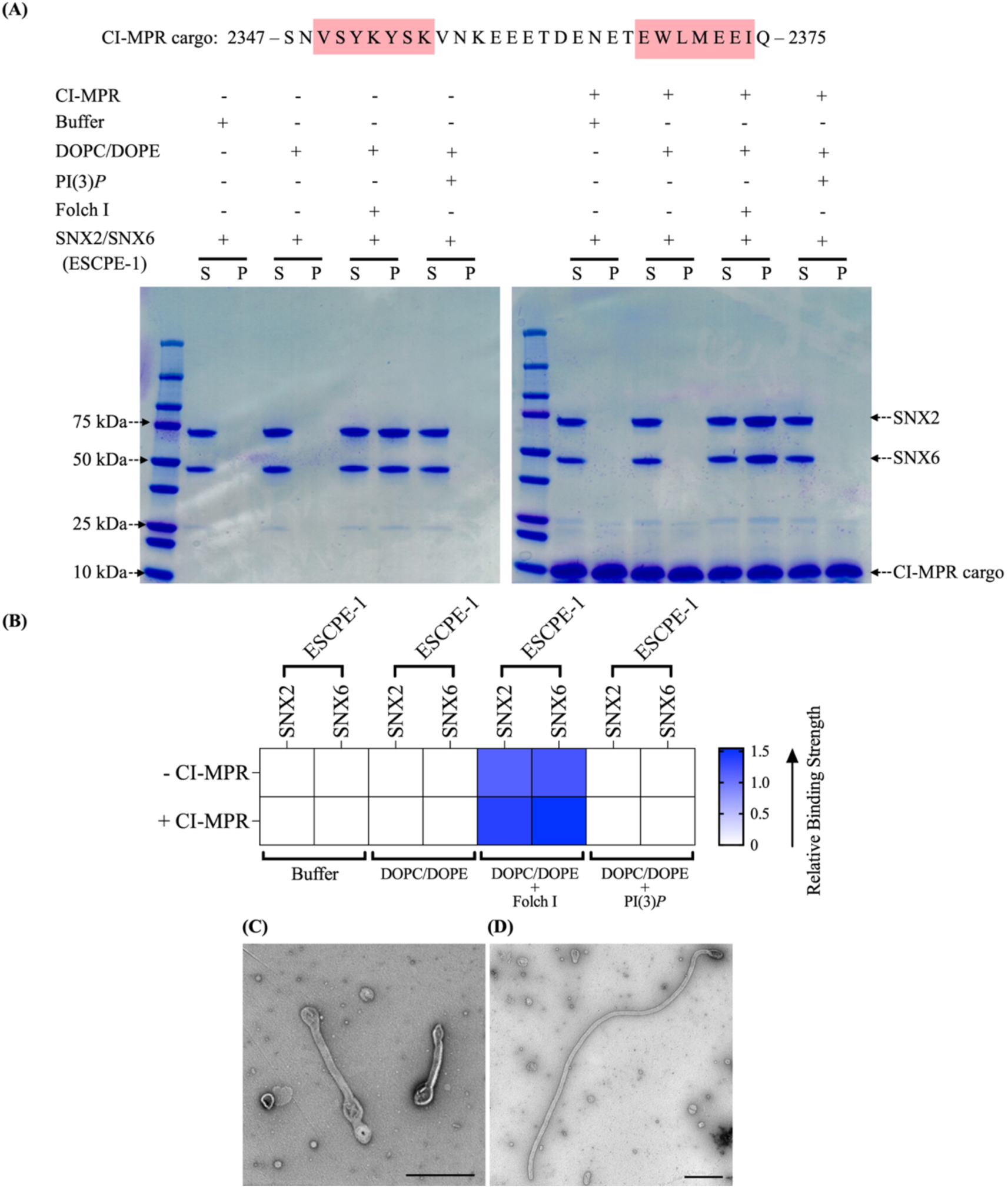
Membrane binding and tubulation properties of ESCPE-1 differ from those of SNX27/ Retromer. (A) Membrane binding of human SNX2/SNX6 (ESCPE-1) complex by liposome pelleting assay. Purified human SNX2/SNX6 complex was incubated with liposomes in the presence or absence of the CI-MPR cargo motif (residues 2347–2375; sequence motifs highlighted in red text). Buffer and DOPC/DOPE alone were used as negative controls to detect non-specific binding. Samples were subjected to ultracentrifugation followed by SDS-PAGE and Coomassie staining of the unbound supernatant (S) and bound pellet (P) fractions. ESCPE-1 is recruited to membranes in the presence of Folch I alone (left gel) and Folch I with CI-MPR cargo (right gel). (B) Binding of ESCPE-1 to phosphoinositide-enriched membranes visualized by SDS-PAGE was quantified by measuring relative protein band intensities (ImageJ) as in Figure 3. (C, D) Negative stain EM reveals tubulation of Folch I-enriched liposomes incubated with SNX2/SNX6 (ESCPE-1) alone (C) or in presence of CI-MPR cargo (D). Scale bars represent 500 nm.

**Figure S9.**
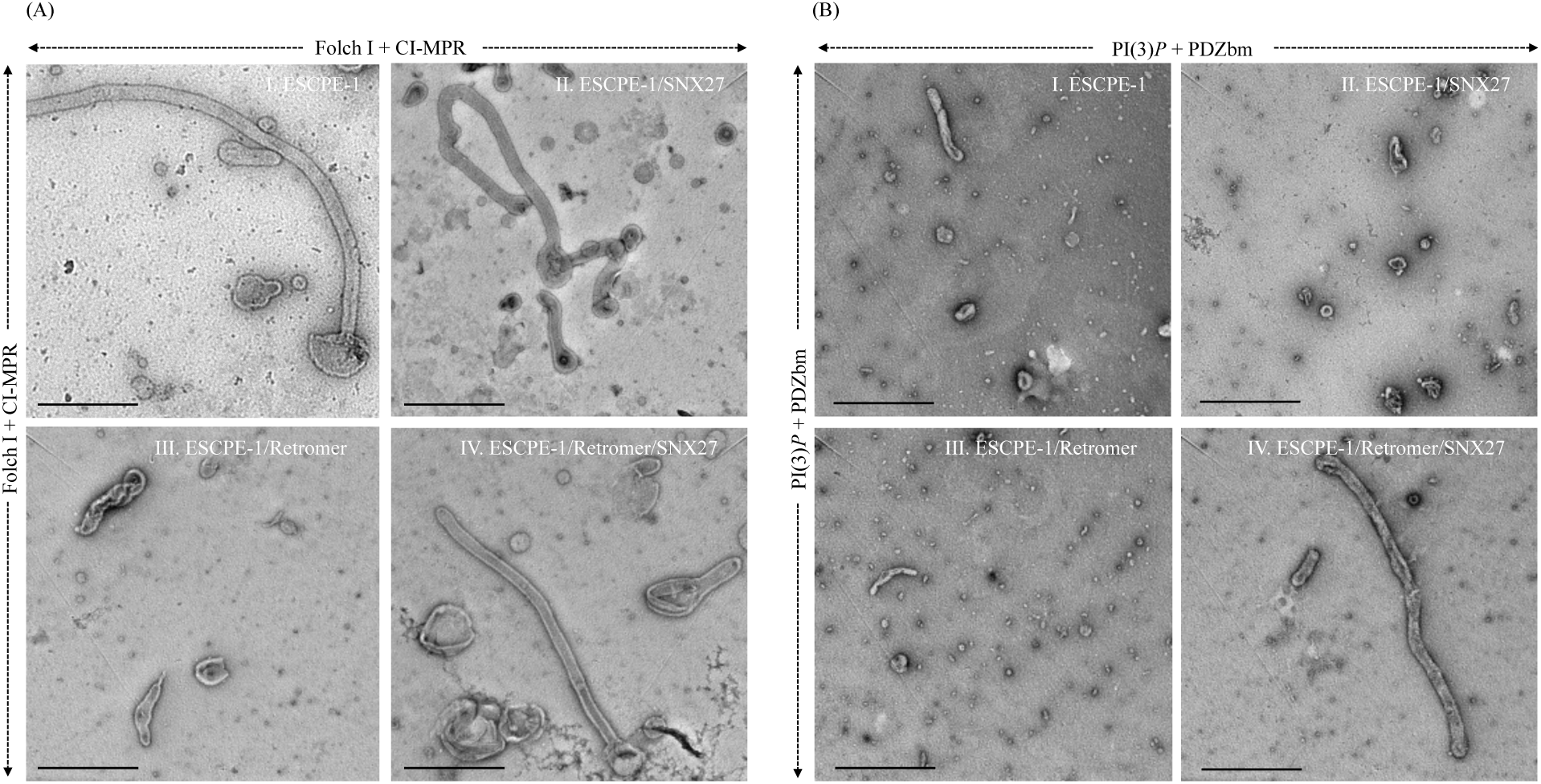
Morphology of membrane tubules generated by endosomal coat proteins visualized using negative stain EM. **(A)** Representative negative stain EM images depicting Folch I-enriched liposomes with CI-MPR cargo motif following incubation with (I) SNX2/SNX6 (ESCPE-1); (II) ESCPE-1 and SNX27; (III) ESCPE-1 and Retromer; and (IV) ESCPE-1, Retromer, and SNX27. These data further suggest ESCPE-1 drives tubulation on its own without contribution from SNX27 or Retromer. SNX27 does not effectively bind Folch-enriched membranes these conditions (see Figure 4A) or contribute to morphology. Retromer does not pellet with ESCPE-1 under these conditions (Figure 4A), and its presence may negatively impact tubule formation (panel III). **(B)** Representative negative stain EM images depicting PI(3)*P*-enriched liposomes with 5-HT4(a)R PDZbm cargo motif following incubation with (I) SNX2/SNX6 (ESCPE-1); (II) ESCPE-1 and SNX27; (III) ESCPE-1 and Retromer; and (IV) ESCPE-1, Retromer, and SNX27. ESCPE-1 is not efficiently recruited to PI3*P*-enriched membranes (see Figure 4B) and probably does not contribute to morphology observed in panel B-IV. Scale bars represent 500 nm.

**Figure S10.**
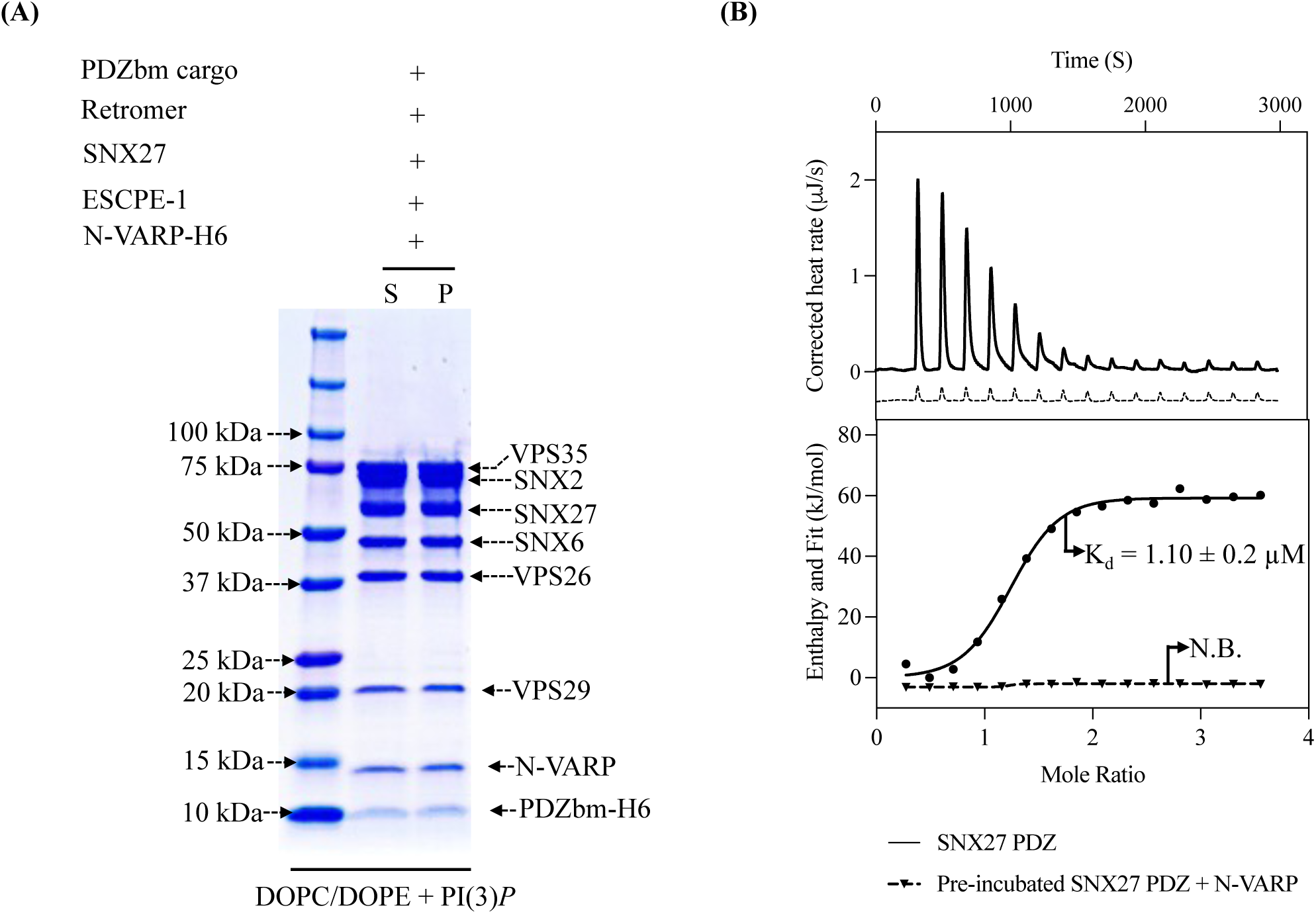
The VARP N-terminus recruits the supercomplex to membranes and binds in SNX27 PDZ cargo binding site. (A) Purified N-VARP protein was incubated with SNX27, ESCPE-1, and Retromer in the presence of PDZbm cargo and PI(3)*P*-enriched liposomes. All protein components are recruited to membranes in the presence of N-VARP alone. (B) Isothermal titration calorimetry (ITC) competition experiments were undertaken to establish whether N-VARP and PDZbm cargo motifs bind the same site on SNX27 PDZ. Synthesized PDZbm cargo peptide from 5-HT4(a)R was titrated into the calorimeter cell containing either purified SNX27 PDZ protein alone (dark black traces) or a 1:1 mixture of purified SNX27 PDZ and N-VARP proteins (dotted black traces). The PDZbm motif binds SNX27 PDZ with a K_D_ near 1 μM as established in the literature, while no detectable binding is observed when PDZbm peptide is titrated into the SNX27 PDZ/N-VARP mixture.

